# Branching topology of the human embryo transcriptome revealed by entropy sort feature weighting

**DOI:** 10.1101/2023.10.12.562031

**Authors:** Arthur Radley, Austin Smith

## Abstract

Single cell transcriptomics (scRNA-seq) transforms our capacity to define cell states and reveal developmental trajectories. Resolution is challenged, however, by high dimensionality and noisy data. Analysis is therefore typically performed after sub-setting to highly variable genes (HVGs). However, existing HVG selection techniques have been found to have poor agreement with one another, and tend to be biased towards highly expressed genes. Entropy sorting provides an alternative mathematical framework for feature subset selection. Here we implement continuous entropy sort feature weighting (cESFW). On synthetic datasets, cESFW outperforms HVG selection in distinguishing cell state specific genes. We apply cESFW to six merged scRNA-seq datasets spanning human early embryo development. Without smoothing or augmenting the raw counts matrices, cESFW generates a high-resolution embedding displaying coherent developmental progression from 8-cell to post-implantation stages, delineating 15 distinct cell states. The embedding highlights sequential lineage decisions during blastocyst development while unsupervised clustering identifies branch point populations. Cells previously claimed to lack a developmental trajectory reside in the first branching region where morula differentiates into Inner Cell Mass (ICM) or Trophectoderm (TE). We quantify the relatedness of pluripotent stem cell cultures to embryo cell types and identify naïve and primed marker genes conserved across culture conditions and the human embryo. Finally, by identifying genes with specifically enriched and dynamic expression during blastocyst formation, we provide markers for staging lineage progression from morula to blastocyst. Together these analyses indicate that cESFW provides the ability to reveal gene expression dynamics in scRNA-seq data that HVG selection can fail to elucidate.

## 1 INTRODUCTION

Single cell RNA sequencing (scRNA-seq) was first described in 2009 (Tang et al. 2009), and has since become a cornerstone of stem cell and developmental biology research. scRNA-seq in principle allows the expression levels of the entire transcriptome of individual cells to be measured in an unbiased manner. The mRNA profiles can then be used as a proxy to define the identity of each cell. A key application is to assess how transcriptomes change as cells transition from one cell state to another and thereby obtain insight into regulatory genes and networks.

Computational tools have been developed to extract useful information from the large and complex datasets that scRNA-seq generates. These bioinformatics pipelines can be broadly broken down into three main steps (Haque et al. 2017):

1. Sequencing alignment - Conversion of the experimentally derived sequencing libraries into an mRNA counts matrix by aligning the detected sequences to a reference genome.
2. Counts matrix processing - Apply quality control and processing techniques to the counts matrix, with the aim of maximising the contribution of biologically relevant expression signals and minimising the presence of technical noise.
3. Data analysis - Combine bioinformatics/data science techniques with domain knowledge to uncover interesting gene expression patterns and dynamics.

There are several well described challenges when working with complex high dimensional datasets such as scRNA-seq data (Lähnemann et al. 2020). Here, we focus on the counts matrix processing stage of scRNA-seq analysis, which is a critical step for increasing ability to extract biologically relevant insights from the data. Due to the importance of counts matrix processing, several open source packages have been released. Notably, Seurat, Scanpy and Scran (Heumos et al. 2023) each provide easy to use workflows for counts matrix processing, which can be further divided into three stages:

1. Basic quality control - Remove low quality cells (low counts per cell, genes per cell, high mitochondrial counts), remove genes expressed in a very low number of cells, etc.
2. Highly variable gene (HVG) selection - Identify a subset of the tens of thousands of genes that are believed to be more informative of cell identity than the rest of the genes.
3. Data denoising - Apply methods such as principal component analysis (PCA) and data regression/smoothing to mitigate the presence of technical artefacts such as stochastic noise or batch effects.

In these conventional workflows, the primary aim of HVG selection is to address the curse of dimensionality (Bellman 1957) by trying to find a subset of the genes in the original scRNA-seq counts matrix that are more discriminative of cell identity. However, poor reproducibility between different HVG methodologies has brought into question the robustness of HVG selection (Yip, Sham, and Wang 2019). Furthermore, HVG selection has been found to be biased towards selecting highly expressed genes over lowly expressed genes.

Following HVG selection, methods that augment or transform the data, such as feature extraction, data smoothing and batch integration are employed to maximise the biologically relevant information that can be extracted (Chu et al. 2022). The prevailing view is that scRNA-seq data has such high levels of inherent experimental noise and/or gene expression stochasticity, that computational methods must be applied to amplify biological signals of interest. While there are many examples where this approach has successfully aided the identification of interesting gene expression profiles, any computational technique that changes/augments the values in the counts matrix prior to downstream analysis has the potential to introduce spurious artefacts. For example, PCA is a feature extraction method that creates a completely new set of features from the gene expression counts matrix, based on a linear transformation of the data. Compression of scRNA-seq data in this manner can lead to undesirable effects, such as combining distinct groups of cells into a single cluster, or removing gradients of gene expression dynamics (Yeung and Ruzzo 2001). Likewise, smoothing techniques that try to impute/repair spurious gene expression values in a scRNA-seq counts matrix based on cell-cell similarities or gene-gene relationships, may introduce computational false negative or false positive expression values that obfuscate real biological signals of interest (Andrews and Hemberg 2018). Similarly, batch correction methods attempt to integrate datasets from different sources by leveraging their commonalities and then changing the values within the datasets so that they become more similar to one another. However, batch correction in this manner risks unintentionally removing heterogeneous biological signals of interest.

The overall aim of HVG selection, feature extraction, data smoothing and batch integration is to try to amplify the biological signal to noise ratio in scRNA-seq data so that downstream analysis can more easily identify gene expression patterns of interest. We hypothesised that it may be possible to obtain a higher resolution of gene expression dynamics in scRNA-seq data through improved feature (gene) selection. We sought to evaluate whether improved feature selection in isolation could increase the biological signal to noise ratio in scRNA-seq data in a manner that mitigates or removes the need for further counts matrix processing via feature extraction, data smoothing or batch integration. A key benefit of reducing the usage of data augmenting processes is that we may be more confident that patterns identified in further downstream analysis are not present due to the introduction of computational artefacts.

We recently outlined a mathematical framework termed Entropy Sorting (ES), and incorporated it into two software packages, Functional Feature Amplification Via Entropy Sorting (FFAVES) and Entropy Sort Feature Weighting (ESFW) (A. Radley et al. 2023). Together FFAVES and ESFW seek to identify genes that are highly structured within the data and hence are more more likely to be predictive of cell identity. Gene selection using these tools exposed the hitherto ambiguous Inner Cell Mass (ICM) population in human pre-implantation embryo data (A. Radley et al. 2023). However, FFAVES and ESFW operate on binarised data such that genes are considered active or inactive in individual cells. Abstracting the data in such a way meant that FFAVES was used to correct sub-optimal gene discretisation, prior to the application of ESFW which assigned feature importance weights. The ESFW weights were then used to select for a subset of all the genes in the data that are more informative of cell identity, in a similar manner to HVG selection. Here, we update ES so that it can be applied to continuous data. We formulate this into a new software package, continuous Entropy Sort Feature Weighting (cESFW) and outline a workflow that utilises cESFW to perform feature selection on scRNA-seq counts matrices. We validate cESFW on synthetic data and then apply the cESFW workflow to human embryo scRNA-seq datasets, achieving high resolution of cell types and developmental trajectories without augmentation or smoothing of the original data. Our analysis provides clear evidence of the existence of the two step model (Cockburn and Rossant 2010) in early human embryo development by elucidating branch points from morula to ICM or TE, and ICM to epiblast (Epi) or hypoblast (Hyp).

These findings demonstrate that cESFW can reveal gene expression states and transitions in scRNA-seq data that were previously unobservable through conventional data analysis techniques.

## 2 RESULTS

### Proposed cESFW feature selection workflow for scRNA-seq data

Data analysis packages such as Seurat/Scanpy/Scran usually involve multiple steps (Fig 1). Many of these steps alter the values in the original counts matrix. For example, HVG selection is highlighted in green because although it reduces the size of the counts matrix by subsetting it down to a smaller set of genes, none of the expression values within the matrix are changed. Conversely, PCA and batch correction methods are highlighted in red because they augment the scRNA-seq counts matrix before passing the data onto the next step of analysis, and hence have a higher potential to introduce spurious computational artefacts.

**Figure 1.**
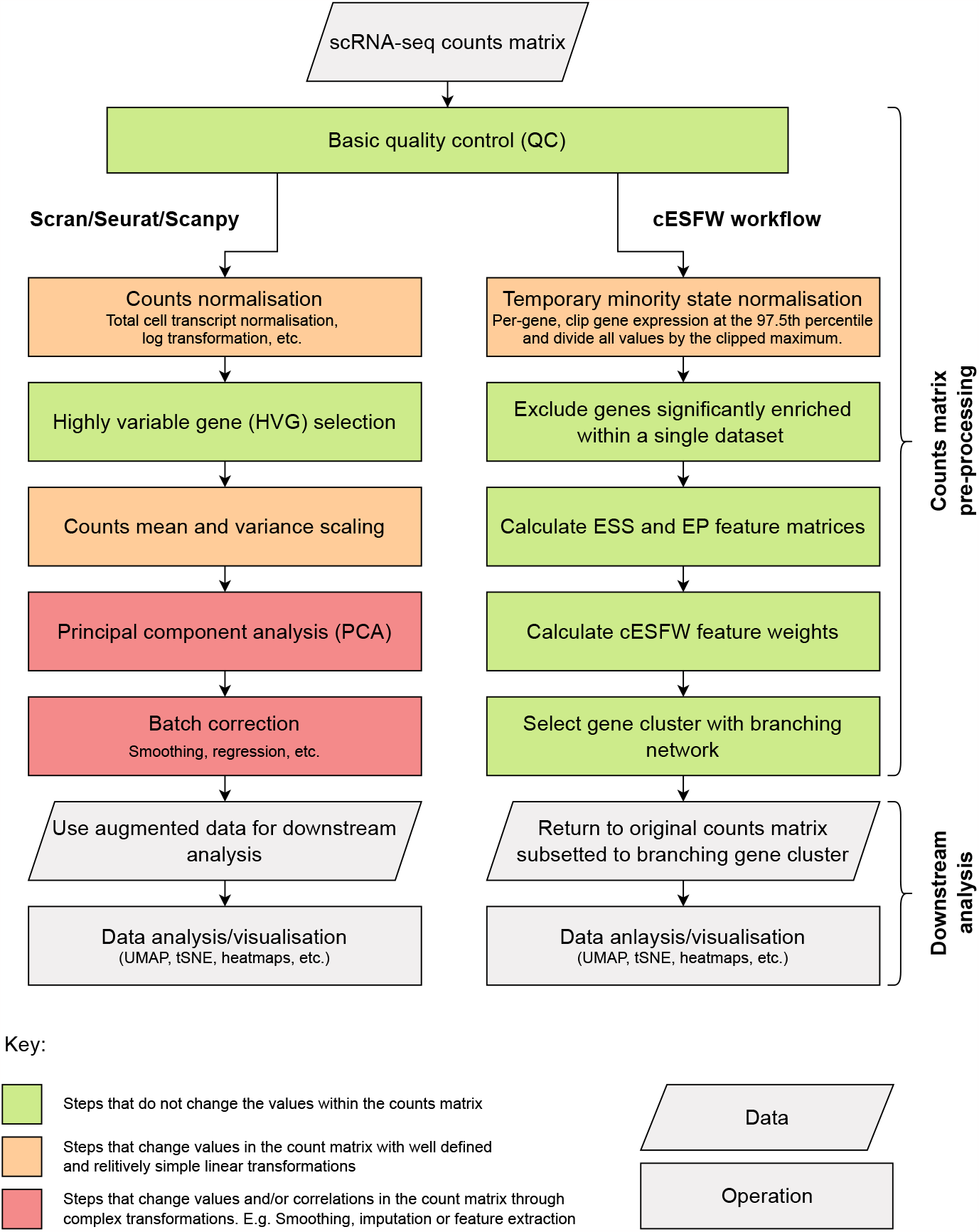
scRNA-seq counts matrix processing workflows. Left hand column shows a high level outline of the workflows implemented by widely used scRNA-seq bioinformatics pipelines such as Scran, Seurat and Scanpy. Right hand column outlines the steps of our cESFW workflow. Colours indicate the degree to which the steps alter expression counts values in the original scRNA-seq counts matrix before the data are passed onto downstream analysis steps.

As an alternative to conventional workflows, we propose cESFW (Fig 1). This workflow facilitates unsupervised gene importance weighting that more robustly identifies genes indicative of cell type than HVG selection. Below we demonstrate that the cESFW gene selection methodology leads to improved downstream analysis, without the need to change any of the expression values in the original counts matrix. This reduces the potential for computational artefacts to be introduced into the analysis of a scRNA-seq dataset compared to Seurat/Scanpy/Scran workflows. A key difference between cESFW and our previous approach, ESFW (A. Radley et al. 2023), is that cESFW can be applied to data with continuous values, whereas ESFW could only be applied to data with binarised values. For a description of how ESFW was adapted into cESFW see SI 1, and for a detailed breakdown of each step of our cESFW workflow, see MATERIALS AND METHODS.

### cESFW robustly discriminates between significant and non-significant gene correlation signals

The foundation of the cESFW workflow is the use of cESFW to perform feature selection in place of HVG feature selection. Both cESFW and HVG feature selection aim to take a high dimensional scRNA-seq dataset and subset it down to a set of genes that are believed to be more informative of cellular identity than the entire gene set. We used synthetic data to perform a side by side analysis of cESFW and the HVG implementations of Scran and Seurat. The Scanpy protocol was excluded because rather than providing a ranked list of genes, Scanpy provides a true/false index for selected HVGs, making it difficult to quantify performance.

We used Dyngen (Cannoodt et al. 2021) to create 4 synthetic datasets (SD) with known ground truths as to which genes relate to cell type specific transcription factor (TF) networks and which are part of ubiquitous housekeeping (HK) gene networks. Dyngen generates synthetic scRNA-seq data by simulating empirically derived gene regulatory networks (GRNs), while allowing control over the size and shape of the generated data. We generated 4 synthetic datasets (Fig 2A). The column labeled “High housekeeping gene expression” refers to a property of Dyngen such that at different random seeds, the average expression of the simulated genes can vary significantly. We took advantage of this to intentionally find random seeds that generated data where either the house keeping (HK) genes had comparable expression levels to the TFs (High housekeeping gene expression = No) or the HK genes had relatively high expression levels compared to the TFs (High housekeeping gene expression = Yes). For example, the average non-zero expression of SD3 TFs and HKs are 9.75 and 1.26 respectively, whereas for SD4 they are 1.74 and 115 respectively. We introduced this layer of analysis because previous evaluations of HVG selection methods have highlighted that the gene selection processes can be biased towards genes that have higher mean expression (Yip, Sham, and Wang 2019). The effect of highly expressed HK genes are shown in Figures 2B and C. In Fig 2B the UMAP generated for SD4 using just the TFs shows the expected branching trajectories, whereas in Fig 2C, including the HK genes for UMAP generation obscures the branching gene expression dynamics.

**Figure 2.**
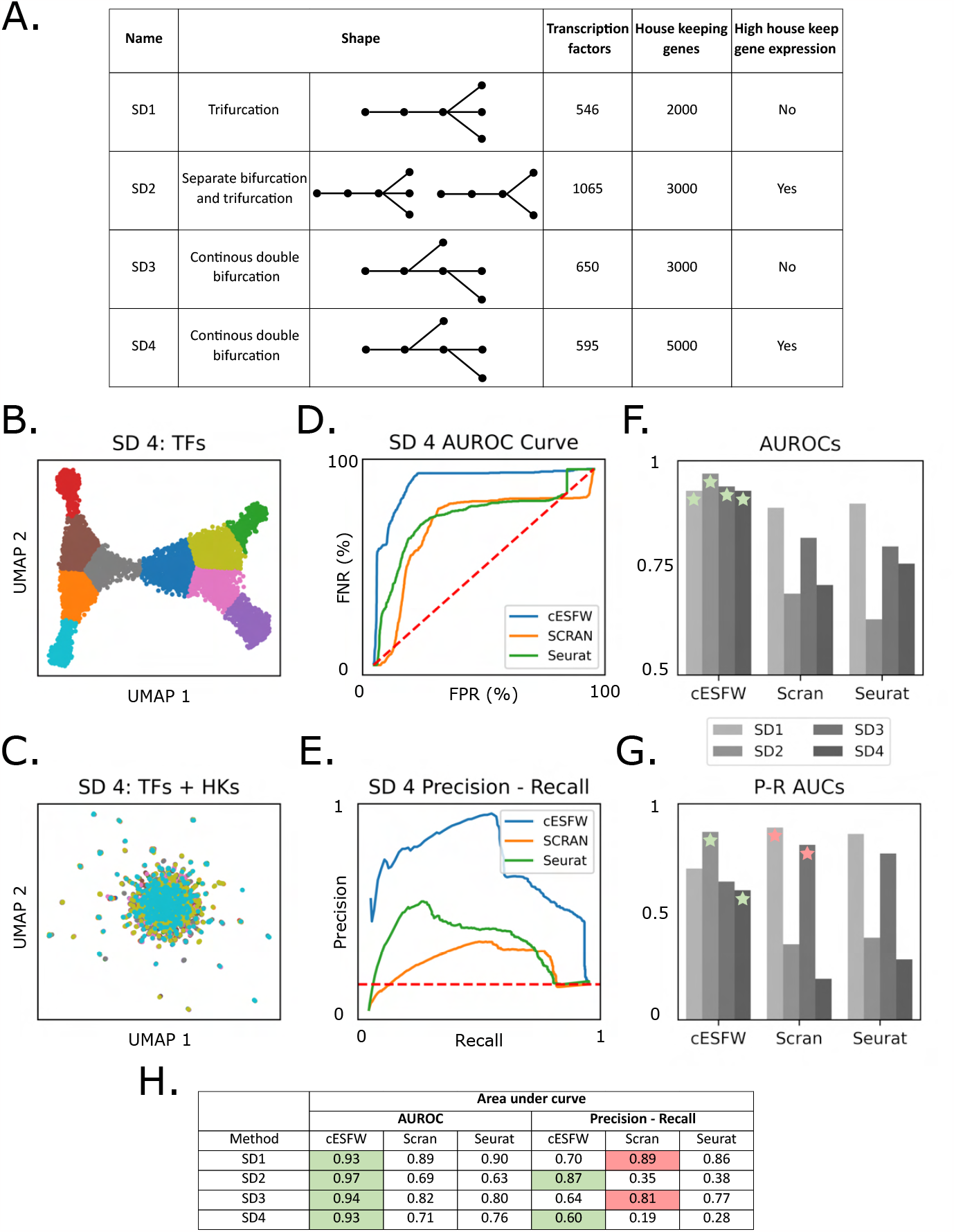
Comparison of cESFW and HVG feature selection techniques. **A.** Summary of the 4 synthetic datasets (SD) of varying size and shape created using the Dyngen software. **B**. UMAP of SD4 created using only the TF genes. **C**. UMAP of SD4 created using all genes (TFs and HKs). **D**. AUROC curve generated using the ranked gene lists from application of cESFW, Scran and Seurat gene selection to SD4. **E**. Precision-Recall curve using the same inputs as D. **F**. AUROC scores after applying cESFW, Scran and Seurat gene selection to all 4 SDs. Green stars indicate that cESFW had the highest score for a SD and red indicate when a different method had a higher score than cESFW. **G**. Precision-Recall AUCs using the same inputs as F. **H**. Summary of all AUC results. Green boxes indicate that cESFW had the highest score for a SD and red indicate when a different method had a higher score than cESFW. For UMAPs and AUC curves relating to SD 1-3, see Fig S1.

Dyngen allows us to differentiate between genes informative of cell type (TFs) and ubiquitously expressed genes (HKs). We used this to quantify the performance of the ranked gene lists generated by cESFW, Scran and Seurat using receiver operating characteristic (ROC) and Precision-Recall (PR) curves (Grau, Grosse, and Keilwagen 2015). For this analysis, the area under (AU) the ROC and PR curves give scores between 0 and 1, where 1 means that the feature selection method is perfect at discriminating between cell type informative TF genes and non-informative HK genes. We show both the ROC and PR curves because ROC is more readily comparable across datasets, whereas PR provides a comparable metric of method performance that is more sensitive to class imbalance (many more HK genes compared to TF genes). Figures 2D and E show the AUROC and PR curves respectively for SD4. In both cases, the area under the blue cESFW curve is largest, indicating that cESFW outperforms Scran or Seurat at ranking TFs as more informative of cell identity than HKs. For SD1 we find that, although cESFW outperforms Seurat and Scran on the AUROC curve, it has the lowest performance on the PR curve (Fig S1C and D). The results of all the ROC and PR curves are summarised in Fig 2F-H. The AUROC scores of cESFW are higher than Scran and Seurat on all 4 synthetic datasets (Fig 2F). High AUROC scores can be thought of as enriching highly ranked genes with genes that are indicative of cell type.

Although cESFW does not have the highest PR-AUCs score for SD 1 and 3 (Fig 2G), we note that the drop in PR-AUCs compared to SD 2 and 4 is relatively small compared to Scran and Seurat. Since SD 2 and 4 are the datasets where HK gene expression is relatively high compared to TF expression, this reinforces previous findings that HVG selection can be biased towards highly expressed genes (Yip, Sham, and Wang 2019). Hence, the results in Fig 2G suggest that cESFW is less sensitive to the presence of highly expressed genes than HVG selection methods.

These results on 4 independent synthetic datasets show that cESFW can provide more robust feature selection than HVG selection.

### cESFW helps reveal high resolution gene expression dynamics in the early human embryo

We next incorporated cESFW into the proposed workflow (Fig 1, right hand column) and applied it to 6 merged independent scRNA-seq datasets of early human embryo development (Yan et al. 2013; Fogarty et al. 2017; Petropoulos et al. 2016; Yanagida et al. 2021; Meistermann et al. 2021; Xiang et al. 2020). These datasets have previously been characterised (Petropoulos et al. 2016; Giuliano G. Stirparo et al. 2018; Meistermann et al. 2021; M. Singh et al. 2023; Wei et al. 2023), and there is a foundation of experimental knowledge defining the expected distinct cell types and stages.

#### Generation of a high resolution day 3-14 human embryo UMAP

The cESFW workflow identified a set of 3012 genes. For details and code see MATERIALS AND METHODS. Gene set enrichment analysis found gene ontology (GO) terms relating to regulation and development (Fig S10B). Using this gene set we obtained a smooth, high resolution UMAP embedding (Fig S2A). Importantly, the UMAP embedding was generated by simply sub-setting down to the 3012 cESFW workflow genes, without any data augmentation or smoothing. The observed structure should therefore reflect biologically significant gene expression patterns, free of computational artefacts that may be introduced through conventional scRNA-seq counts matrix processing workflows (Fig 1, left hand column).

The UMAP clearly displays continuous progression along the developmental time course. Cells from the 5 pre-implantation embryo datasets are intermingled across the UMAP space, indicating minimal contribution of batch effects (Fig S2B). Cells from the post-implantation extended culture samples (Xiang et al. 2020) form groups adjacent to distinct regions of pre-implantation cells, indicating that related gene expression patterns have been identified across the pre- and post-implantation datasets.

We performed unsupervised agglomerative clustering on the scRNA-seq counts matrix subset down to the 3012 cESFW genes. Overlaying these clusters onto the UMAP embedding (Fig S2A) provides an unsupervised foundation to label samples with 15 distinct cell type/stage annotations (Fig 3C, Fig S2B, C). Comparison with annotations from previous studies (Petropoulos et al. 2016; Giuliano G. Stirparo et al. 2018) corroborated our annotations and UMAP embedding (Fig S2D, E). Notably, the unsupervised clusters were readily annotated, whereas previous annotations were generated through supervised analysis focused on a handful of candidate marker genes. A gene expression heatmap of the cESFW genes grouped by our 15 cell type annotations shows cell type specific gene expression profiles (Fig 3D). Genes implicated in the literature to correspond with 13 of our cell type annotations are represented in the 3012 cESFW gene set (Cockburn and Rossant 2010; Taubenschmid-Stowers et al. 2022; Giuliano G. Stirparo et al. 2018; M. Singh et al. 2023; Corujo-Simon, A. H. Radley, and Nichols 2023; Liu et al. 2022; Zadora et al. 2017; Yue et al. 2020; Yabe et al. 2016; Yang et al. 2021). Such genes show specific gene expression in the UMAP embedding (Fig 3E, Fig 5A Fig S3). Mural trophectoderm (TE) lacks validated specific markers but is known to express HAND1 (Liu et al. 2022). Although HAND1 is not captured in the cESFW gene set, it shows relatively specific day 6/7 Mural TE expression on our UMAP embedding Fig S3).

**Figure 3.**
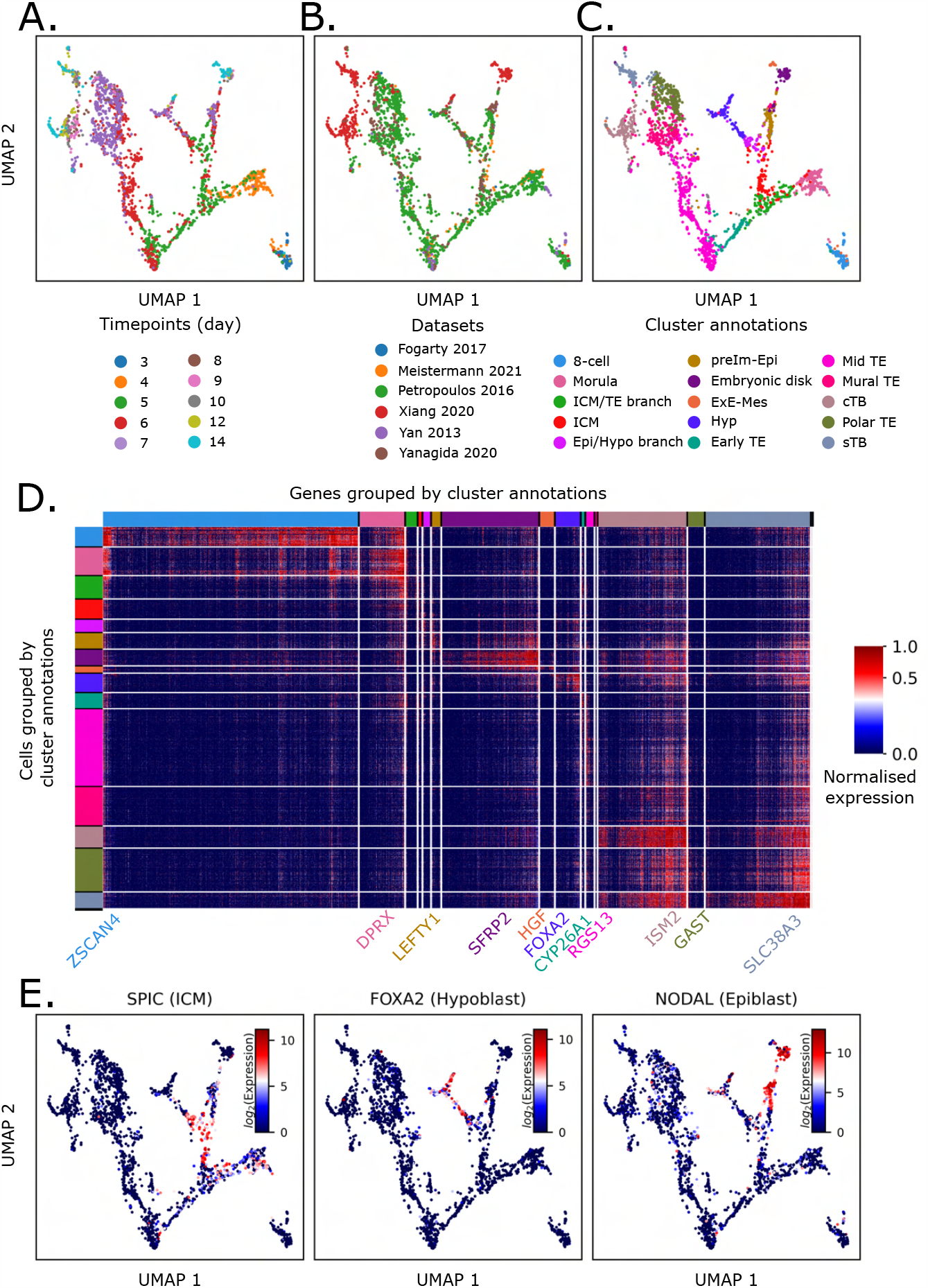
High-resolution UMAP embedding of early human embryo development. **A-C**. UMAP embedding generated using the 3012 genes identified by our cESFW workflow, **A**. coloured by time point labels from data source papers, **B**. coloured by dataset sources, **C**. coloured by our cell type cluster annotations. preIm-Epi = pre-implantation epiblast, ExE-Mes = extraembryonic mesenchyme, Hyp = hypoblast, cTB = cytotrophoblast, sTB = syncytiotrophoblast. See Fig S2 and Fig S3 for further details regarding the cluster annotations. **D**. Heatmap of the 3012 cESFW selected genes, grouped by our cell cluster annotations. **E**. Known cell type marker expression profiles validate our UMAP annotations. See Fig S3 for more examples.

Visualisation of chronological embryo progression with gene expression patterns consistent with the literature demonstrates that the cESFW workflow identifies a set of genes that is highly descriptive of the transcriptional landscape of day 3-14 developing human early embryos.

#### Feature selection via cESFW outperforms HVG selection

To compare the performance of cESFW with the popular Scran and Seurat workflows using the same input data, we identified the 3012 top ranked genes for each technique.

Previous studies comparing HVG selection methods have shown that there is often poor concordance between gene sets selected by different methods (Yip, Sham, and Wang 2019). Intersecting the sets of 3012 genes from the cESFW, Scran and Seurat workflows, we find only 290/3012 (9.63%) of the genes are common to all 3 gene sets (Fig S4A).

UMAPs generated using the Scran and Seurat HVGs were able to broadly separate samples by their embryo time points, but did not provide a smooth embedding with a coherent progression of cell types and developmental trajectories (Fig S4C, D).

We used silhouette scores to quantify how well each of the 3012 gene sets formed distinct clusters of samples, using our UMAP cell type annotations (Fig 3C) as cluster labels. Silhouette scores measure how similar samples are to their designated clusters, with positive scores indicating samples are most similar to the cluster they are a member of, and negative scores designating samples that would be better assigned to a different cluster. The average of the silhouette scores for all samples within a cluster then quantifies the quality of the cluster as whole. The cESFW genes produced higher silhouette scores than the Seurat or Scran gene sets for 14 out of the 15 clusters with similar scores for ICM (Fig S4B). Notably the Epi/Hyp branch, Hyp and Mid TE score positive with cESFW but have negative silhouette scores using HVG, signifying that the HVG gene sets are unable to identify these cell types as distinct from other cells in the data.

These results show qualitatively and quantitatively that cESFW gene selection outperforms popular HVG selection workflows on these human embryo scRNA-seq data and help explain why the UMAP generated using the 3012 cESFW gene set is able to readily separate known cell types from one another.

### Lineage branching blastocyst development

The textbook model of blastocyst formation derived from studies in the mouse embryo entails sequential lineage bifurcations that generate first trophectoderm (TE) and inner cell mass (ICM), and then hypoblast (Hyp) and epiblast (Epi) (Cockburn and Rossant 2010). However, the existence of two branch points in human embryogenesis has been questioned in previous analyses of scRNA-seq data (A. Radley et al. 2023). More recent studies have proposed two bifurcations, but have not demonstrated separate branch point populations (Yanagida et al. 2021; A. Radley et al. 2023; Wei et al. 2023).

The cESFW embedding unambiguously reveals two binary branch points in pre-implantation embryogenesis; from morula to ICM or early TE, and from ICM to Epi or Hyp.

Furthermore, unsupervised clustering identifies populations of cells at these intersections which we labeled ICM/TE and Epi/Hyp branch cells, respectively (Fig 3A). Silhouette scores quantify both as distinct groups of cells according to transcriptional profile (Fig S4B). To examine the inferred lineage trajectories we performed RNA velocity analysis (Bergen et al. 2020) (Fig 4D). The RNA velocity vectors are aligned well with temporal ordering and lineage divergence across the embedding (Fig 4B). Notably, RNA velocities show a progression of morula samples towards the ICM/TE branch cells, which then diverge either towards the ICM or to early TE.

**Figure 4.**
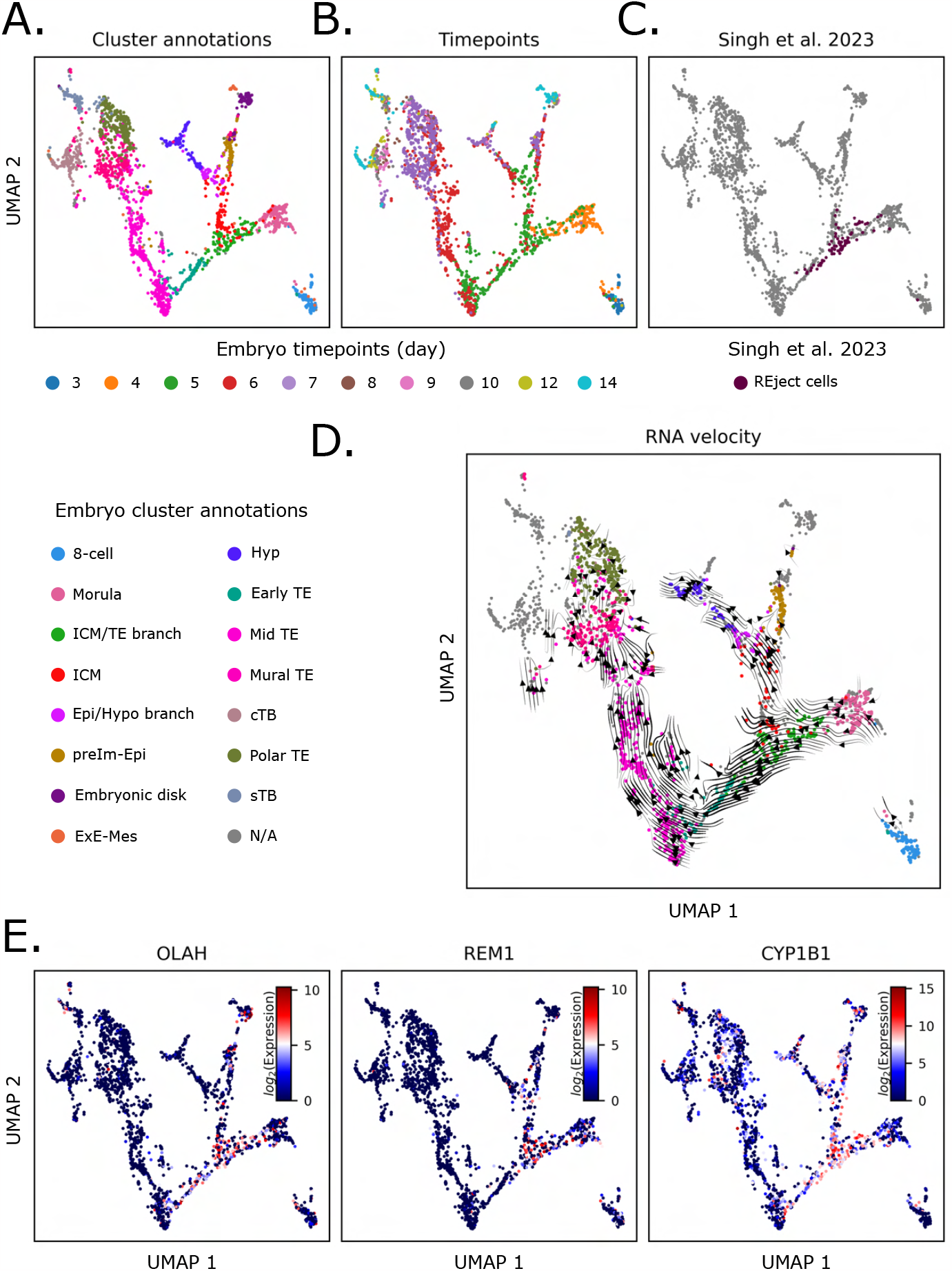
REject cells are within the ICM/TE branch point cluster. UMAP embedding generated using the 3012 genes identified by our cESFW workflow, **A**. coloured by time point labels from data source papers, **B**. coloured by dataset sources, **C**. coloured by M. Singh et al. 2023‘s NCC sample labels. **D**. RNA velocity vectors overlaid onto the UMAP embedding. Samples in grey were not part of the RNA velocity analysis owing to the scRNA-seq spliced and unspliced counts matrices only being available for the Petropoulos et al. 2016 dataset. **E**. Proposed ICM/TE branch marker genes.

In Fig 4E, we use our UMAP and ranked gene list (Table S1) to highlight 3 genes that show relatively specific upregulation within the ICM/TE branch population.

In a recent publication, M. Singh et al. 2023 re-analysed the Petropoulos et al. 2016 human embryo scRNA-seq dataset and found a distinct and previously undescribed group of day 4/5 cells. However these authors were unable to place this cluster in a chronological differentiation trajectory and noted upregulation of several proposed apoptosis markers, leading them to hypothesise that this group of cells had accumulated DNA damage and was fated to apoptose. They therefore refer to them as “REject” cells. However, when we inspected the REject cells in the cEFSW embedding we found that they do not constitute a separate cluster, but were almost entirely within the ICM/TE branch (Fig 4C). We therefore investigated the markers suggested to indicate that the REject cells are pre-apoptotic. We found that most of these genes are expressed equally highly elsewhere in the developing human embryo, suggesting either that they are not strong discriminators of pre-apoptotic versus non-apoptotic cells, or that apoptosis is distributed across cell types (Fig S5). For example, although suggested apoptosis markers BIK, ATG2A and ATF3 are up-regulated in the ICM/TE branch when compared against the early TE, ICM, preIm-Epi and Hyp populations, these 3 genes are more highly expressed in the 8-cell and/or morula populations (Fig S5B). In a similar vein, BAK1, CTSB and CASP6 were suggested as apoptotic markers specific to the REject population, but we find these genes to be broadly expressed across several cell types of day 3-14 human embryos (Fig S5C). Thus, we do not find strong indications that the proposed REject cells are specifically fated for apoptosis.

Overall, these findings substantiate the occurrence of an initial lineage bifurcation between TE and ICM on day 4/day 5 that precedes segregation of ICM to hypoblast and epiblast between day 5 and day 6.

### Comparison of human pluripotent stem cell cultures to reference human embryo scRNA-seq embedding

Human pluripotent stem cells (PSCs) are generally considered to be analogues of the pluripotent lineage in the embryo, the epiblast. However, PSCs exist in different states that are propagated in distinct signaling environments (Smith 2017; Nichols and Smith 2009). Correlation of PSCs with developmental stages of epiblast in the human embryo has been problematic due to the scarcity of relevant data, and the limited resolution provided by previous analysis methods. We took the opportunity provided by our high-resolution embedding to re-evaluate the transcriptome relatedness of various cultures of human PSCs to cell types in the day 3-14 embryo, highlighting pre- and post-implantation epiblast (Fig 5A, Fig S3).

**Figure 5.**
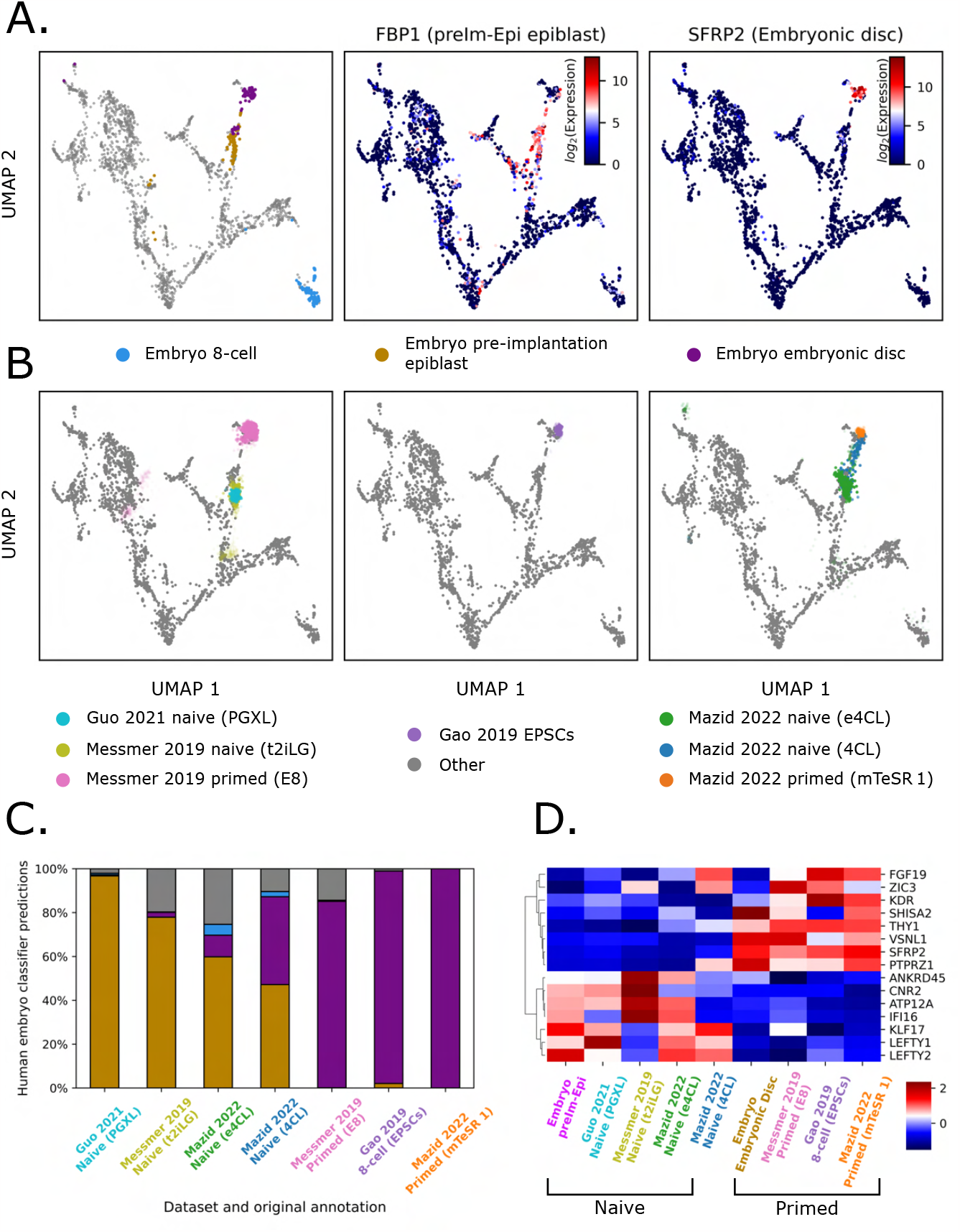
Characterising the human embryo counterparts of *in vitro* stem cell cultures. **A.** UMAP embedding highlighting the embryo 8-cell, preIm-Epi and Embryonic disc populations. FBP1 and SFRP2 shown as previously identified preIm-Epi and Embryonic disc markers respectively (Messmer et al. 2019). **B**. Projection of cultured naïve, primed and EPSC scRNA-seq datasets onto our UMAP embedding. To aid interpretation, sample transparencies are scaled by the density of samples on the embedding from the same condition, such that samples in the densest region are completely opaque. Abbreviations in brackets denote relevant culture media. **C**. Quantification of cultured naïve, primed and EPSC scRNA-seq sample identities. **D**. Heatmap of z-score normalised gene expression of 8 naïve specific and 8 primed specific gene markers. All values are pseudo-bulk expression values, generated by grouping single cells by their dataset and predicted cell type labels from panel C., and taking the average expression of each group.

We examined published scRNA-seq datasets from conventional primed PSCs and cells in two naïve culture conditions (PXGL (Guo, Giuliano Giuseppe Stirparo, et al. 2021) and its predecessor t2iLGö (Messmer et al. 2019)). We also assessed data for expanded potential stem cells (EPSCs), originally suggested to be related to the 8-cell stage (Gao et al. 2019). We used the *umap*.*transform* function of the Python UMAP package (McInnes et al. 2018) to map the query cells from the PSC cultures onto our reference human embryo UMAP. This method places individual query samples closest to their most transcriptionally similar counterparts in the UMAP in an unsupervised manner based on all 3012 cESFW genes. Figure 5B shows the projections of individual cells from the query scRNA-seq datasets onto the embedding. Conventional primed PSCs are placed in the vicinity of the d10-14 embryonic disk epiblast. EPSCs are similarly positioned close to post-implantation epiblast with *<* 2% of cells adjacent to earlier stages. These findings are consistent with previous comparative analyses with non-human primate embryos (Guo, Giuliano Giuseppe Stirparo, et al. 2021; Nakamura et al. 2017). In contrast, naïve PSCs overlie pre-implantation epiblast (d6-7). Notably cells in PXGL (Guo, Meyenn, et al. 2017) are more homogeneous than cells in t2iLGö (Takashima et al. 2014). Quantification using a k-nearest neighbour classifier trained on the embryo cell type annotations (Fig 5C) assigns 97% of PXGL cells to pre-implantation epiblast compared with 77% for t2iLGö.

We also evaluated published data for recently reported 4CL and e4CL cells that are proposed to be related to d5 ICM and 8-cell stages (Mazid et al. 2022). These samples are more heterogeneous than other PSC cultures. They comprise pre-implantation epiblast-like proportions of 48% for 4CL and 60% for e4CL. A substantial fraction (39%) of 4CL cells are post-implantation epiblast-like while many e4CL cells are positioned between pre- and post-implantation epiblast, indicating either an indeterminate mixed identity or potentially transitioning between states. e4CL cultures contain only 5% of cells which are related to d5 ICM and 5% of cells related to the 8-cell embryo.

#### Identifying naïve and primed hPSC markers

Previous studies have identified gene markers for naïve and primed hPSCs by examining differential expression between these two PSC types specifically. We sought to characterise markers in the context of the entire day 3-14 human embryo. We started by taking the top 3 naïve versus primed PSC markers identified by Messmer et al. 2019 and plotted them on our UMAP embedding (Fig S6). We saw that although these 6 markers were clearly up/down-regulated between the preImp-Epi and embryonic disc samples, for 5 out of the 6 genes the up/down-regulation is not restricted to epiblast lineage. In contrast, the top 3 ranked genes for our UMAP annotated preImp-Epi and embryonic disc cell types (Table S1, Fig 3C), show more restricted expression to the preImp-Epi or embryonic disc regions (Fig S6).

We then proceeded to use our rank gene lists to identify genes that showed specific up-regulation in the preImp-Epi or embryonic disc, while also showing differential expression in naïve versus conventional hPSCs. In Fig 5D we present a heatmap of 8 naïve and 8 primed marker genes that have consistent cell type specific differential expression across the cell line datasets, and in our reference human embryo embedding. We also note that in this heatmap, the 4CL cells show upregulation of both naïve and primed markers, consistent with our previous observation that this culture condition displays mixed identity (Fig 5B, C).

This analysis of naïve, primed and EPSC stem cells from different research groups demonstrates how access to an unbiased view of the early human embryo can facilitate classification of in vitro cell cultures. Furthermore, we show that taking a more holistic approach to cell type marker identification can provide more specific cell type marker genes.

### Emergence of epiblast, hypoblast and trophecto-derm signatures during blastocyst formation

Between days 3 and 7 post fertilisation, the morula of the human embryo develops into the blastocyst, comprising three distinct lineages, Epi, Hyp and TE. We sought to use the cESFW embedding to identify genes that may be useful markers for staging cell identities during blastocyst formation.

The Entropy Sort Score (ESS), is a correlation metric derived from Entropy Sorting (A. Radley et al. 2023). A potential benefit of the ESS for gene ranking compared to other typical methods such as the t-test or Wilcoxon test is that the ESS was specifically derived to reduce whenever gene expression is observed outside of a specific population of interest. Therefore, when there are multiple cell types within a dataset, the ESS can be better suited for identifying regions of specific gene expression, as opposed to more general up/down regulation. Using the ESS we rank genes by how well their expression profiles overlap with the Epi, Hyp or TE morula to blastocyst trajectories, while also progressively restricting the analysis to the terminal Epi, Hyp or TE populations. For examples and details regarding this progressive restriction, see Fig S7 and EXPERIMENTAL PROCEDURES.

In Figures 6A-C we present heatmaps highlighting sets of genes that show sequential up-regulation as the morula differentiates into Epi (A), Hyp (B) or TE (C). Day 7 TE comprises both mural and polar TE. Figures 6D-F and Fig S7 show example genes with specific up-regulation along each trajectory. For example during the emergence of TE cells during blastocyst formation, SLC28A3 (Figures 6F) is first upregualted in the morula and maintains expression until day 7 TE, whilst being downregulated in the ICM, Epi and Hyp lineages. Conversely, the TE marker HAPLN1 (Fig S7B) is upregulated later than SLC28A3 at around day 6 and is maintained in day 7 of TE.

**Figure 6.**
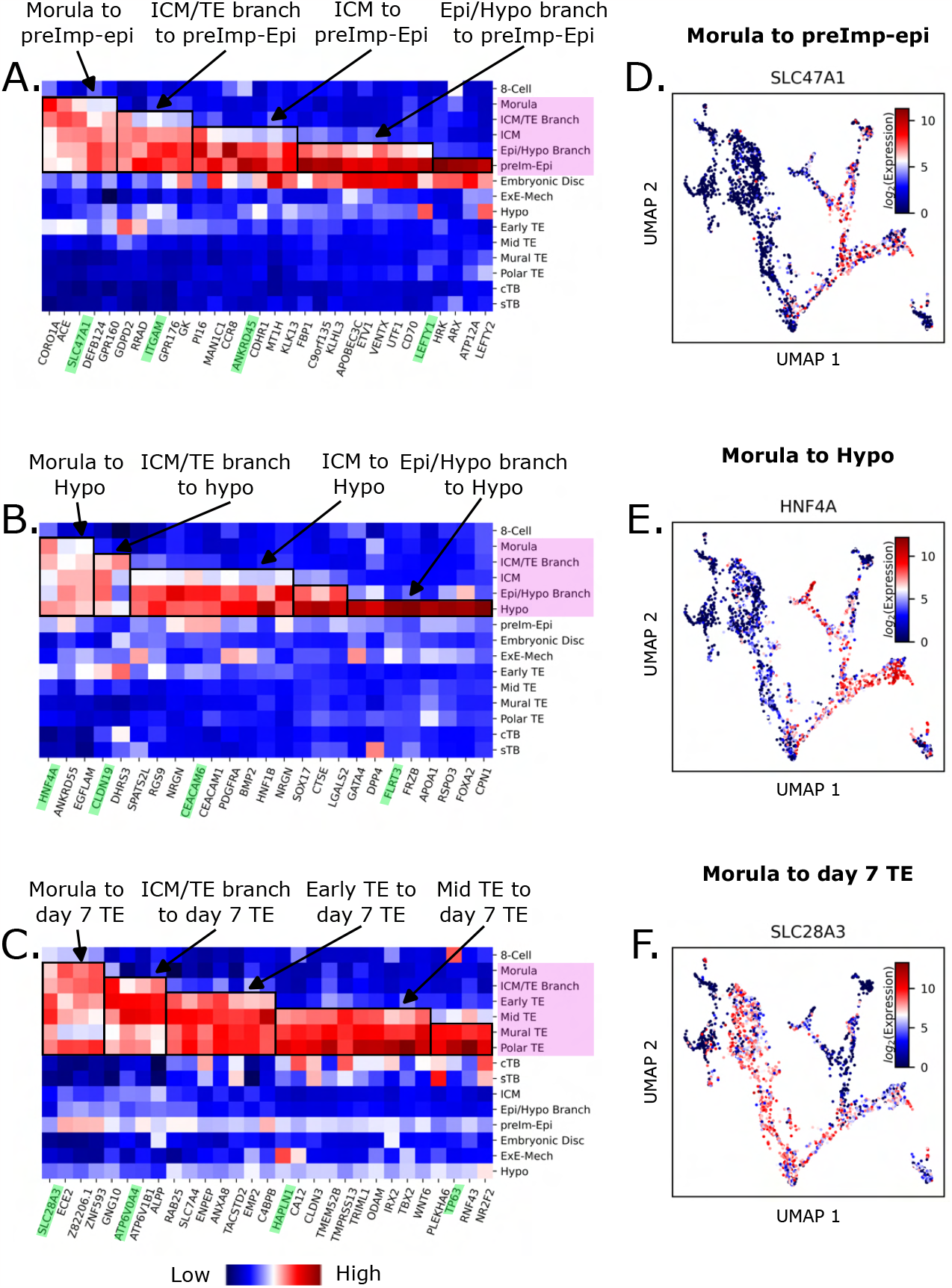
Emerging epiblast, hypoblast and trophectoderm signatures during blastocyst development. **A-C.** Heatmaps of pseudobulk, z-score normalised human embryo data, displaying genes that show blocks of sequential up-regulation between morula and Epi (A.), Hyp (B.) or TE (C.) lineages. Pink boxes highlight most relevant cell types for each lineage. Green highlighted genes are presented in UMAPs within this figure and in Fig S7. **D-E**. UMAP gene expression profiles for example genes showing expression from morula to Epi (D.), Hyp (E.) or TE (F.) lineages.

These findings illustrate how ES and our UMAP embedding can be used to identify new gene expression patterns that may aid future experiments and analyses.

## 3 DISCUSSION

In this work, we introduced cESFW, a feature selection method designed for distinguishing between cell state informative and uninformative genes in high dimensional scRNA-seq data. A key ethos of the cESFW workflow is that the final scRNA-seq counts matrix does not have its values changed in any way. We show that using cESFW to subset down to a set of informative genes is not only sufficient to identify dynamic gene expression states, but can outperform more complex methods that augment the values of the counts matrix. These findings challenge the view that scRNA-seq data is inherently too noisy to identify high resolution gene expression dynamics without repairing, smoothing, or transforming the data first.

On synthetic scRNA-seq data we found that cESFW performs comparably or better than conventional HVG selection. Whereas HVG selection is particularly sensitive to the presence of highly expressed genes (Yip, Sham, and Wang 2019), which lead to a significant drop in ability to distinguish between cell type specific genes and housekeeping genes, cESFW is relatively agnostic to the presence of genes with high mean expression, with a minimal drop in performance. Less influence of expression level when ranking gene importance may be a significant contributor to cESFW’s superior performance on biological data, since evidence is lacking that more highly expressed genes are generally more discriminatory of cellular identity. In addition, the multivariate approach of cESFW may be more robust than univariate HVG feature selection. In the univariate approach, genes are considered in isolation, leading to their importance ranking being more sensitive to fluctuations in properties such as mean expression. cESFW, in contrast, considers pairwise the expression profile of an inspected gene with every other gene in the data, and generates a gene importance weight by quantifying the degree to which the inspected gene forms networks of significantly co-expressing genes.

Applying our cESFW workflow to early human embryo scRNA-seq data generates a high resolution UMAP embedding of different cell types, trajectories and branch points in the developing embryo. Our cell type annotations formed quantifiably more distinct clusters when using top rank genes selected by cESFW compared to the top ranked genes from conventional HVG selection. Furthermore, we observe coherent dataset integration without having to apply batch correction methods as in previous analyses (Meistermann et al. 2021; Wei et al. 2023). We propose that cESFW achieves good integration of datasets from different sources because improved feature selection tips the biological signal to noise ratio in favour of the biologically relevant expression profiles in the data. Thus, cell similarity metrics may be less affected by confounding gene expression signals

Our high resolution map of the early human embryo transcriptome allowed us to substantiate the existence of two distinct branching populations, at the intersections of morula with ICM and TE, and of ICM with epiblast and hypoblast. Their identification and the general topology of the cESFW embedding indicate that human embryogenesis proceeds via the sequence of lineage bifurcations demonstrated in the mouse embryo (Cockburn and Rossant 2010). The branching populations reside at critical junctures in blastocyst formation, the partitioning of extraembryonic and embryonic lineages. Cells in these clusters become specified to alternative fates but may not be committed. For example, PDGFRA has been implicated as a hypoblast marker (Corujo-Simon, A. H. Radley, and Nichols 2023), whereas NANOG has been demonstrated as an epiblast marker (Allègre et al. 2022), yet both markers are heterogeneous in Epi/Hyp branching population. Notably the cluster boundaries extend beyond the topological branch point, which may indicate that cells remain plastic and could be respecified. Further analytical and experimental studies may focus on heterogeneity and gene expression dynamics in these populations to expose the process of lineage choice and commitment.

We also noted that the ICM/TE branch population includes cells previously unannotated and therefore conjectured to be rejected from embryo development. Those cells do express various markers of a pre-apoptotic state, but so do other cells throughout the data. Quantitative immunostaining analyses are needed to reveal if there is a consistently higher frequency of in the embryo.

We used the cESFW gene set and embedding as a reference point to evaluate the identity of various hPSC cultures. Quantifying proportional relatedness of cell lines to embryo populations, confirmed that conventional hPSCs and EPSCs are most related to post-implantation embryonic disc while naïve PSCs are similar to pre-implantation epiblast. This analysis also demonstrated that PXGL naïve PSCs have a very high (97%) representation of pre-implantation epiblast-like cells.

Finally, we used ESSs and our UMAP embedding to suggest sets of genes that may aid cell type staging during days 3-7 of lineage progression during human blastocyst formation. Immunostaining will be of interest to assess protein expression dynamics for these genes.

To conclude, this study demonstrates that cESFW, an algorithm derived from the mathematical framework of ES, has the ability to reveal gene expression dynamics in scRNA-seq data that were previously unobservable. Our characterisation of early human embryo scRNA-seq data has provided ranked gene lists for multiple cell types and a high resolution UMAP embedding. We provide a proposed workflow for incorporating cESFW into general scRNA-seq analysis. We envisage that the cESFW technique and workflow will readily be adapted and improved for application to datasets of varying complexity and type, such as multi-omics or spatial transcriptomics studies.

## 4 Limitations of the study

The benefits of improved feature selection through cESFW, do not preclude the use of other data analysis techniques such as batch integration and data repair. We suggest that more accurate feature selection has the potential to further improve results of all downstream steps. We anticipate that researchers will create adaptations of our proposed cESFW workflow that incorporate a range of data analysis methodologies.

We expect that as more embryo scRNA-seq datasets become publicly available, re-analysis should yield greater resolution. A higher density of samples from day 4-6 embryos will be particular value for better delineation of ICM progression lineage branch points. Further data from extended cultures will enable greater resolution of embryonic disc progression and emergence of extraembryonic mesenchyme and amnion.

The genes we have highlighted for staging of cells during blastocyst formation were picked through semi-supervised analysis. The initial gene sets were curated from ranked gene lists output based on the ESS. However, the final sets of selected genes were subject to manual inspection of the top 30 ESS ranked genes, and genes that appeared to be poorly expressed in the cell populations of interest were manually excluded. Therefore the presented gene sets may be incomplete.

## 5 EXPERIMENTAL PROCEDURES

### Data and code availability

Instructions to install cESFW can be found at, https://github.com/aradley/cESFW.

Computational workflows and data used for the generation of results in this article may be found at https://github.com/aradley/cESFW_Embryo_Topology_Paper.

A copy of the human pre and post implantation embryo data used in this work may be found in the following permanent Mendeley Data repository https://data.mendeley.com/datasets/34td4ds2r9/draft?a=c464ae1c-08a6-430a-8b6dc8146d61f5. (**Please note that until this work is peer reviewed, this Mendeley Data repository link will be in preview mode, and may be subject to change**.)

The data used to create our the UMAP embedding are a combination of human embryo pre-implantation and postimplantation embryo (extended culture) scRNA-seq data. The pre-implantation raw counts scRNA-seq data from Yan et al. 2013, Petropoulos et al. 2016, Fogarty et al. 2017, and Meistermann et al. 2021, were compiled into a single gene expression matrix kindly provided by (Meistermann et al. 2021). The Yanagida et al. 2021 human pre-implantation embryo data are available via GEO accession number GSE171820. The 3D cultured post-implantation human embryo raw counts scRNA-seq data are from Xiang et al. 2020, available at GSE136447. For our analysis, we removed Xiang et al. 2020‘s day 6 and 7 cultured embryo samples to mitigate the contribution of batch effects and uncertainty over timings of embryo stages.

### Synthetic data generation

The 4 synthetic datasets used in this study were generated using the Dyngen software (Cannoodt et al. 2021). Synthetic dataset size and shapes were all controlled using out of box Dyngen functions.

### UMAPs

For consistency, or all UMAPs presented in this work were created with the Python *umap-learn* package, with *n_neighbors* and *min_dist* set to 50 and 0.1 respectively. The correlation distance metric was used instead of the default euclidean distance metric.

All UMAPs displaying gene expression profiles show the *log*_2_ normalised gene expression. No data smoothing or clipping were applied.

### Unsupervised clustering

Unsupervised clustering of scRNA-seq embryo data (Fig S2A) was performed using the Python package, *sklearn*.*cluster. AgglomerativeClustering*, with *k* = 30 clusters and the correlation distance metric.

### Gene marker identification

Identification of markers for UMAP plotting and our ranked gene list (Table S1) was performed by using the Entropy Sort Score (ESS) correlation metric to rank genes that are enriched to a cell type or cell types of interest. For details see A. Radley et al. 2023, and for examples see the workflows provided alongside this paper.

### RNA velocity

RNA velocity was carried out using the scVelo (Bergen et al. 2020) with default parameters, expect for the cell nearest neighbour matrix, which we manually substituted with the cell nearest neighbour matrix according to the 3012 cESFW genes and the correlation distance metric.

### Query cell classification

Query scRNA-seq samples were transformed onto our UMAP embedding using the out of box *umap*.*transform* function. After query samples are placed on the UMAP embedding, we may predict their cell type in the low dimensional UMAP space.

To predict cell identity, we used the *sklearn*.*neighbors. KNeighborsClassifier* Python function. A knn classifier was trained using our UMAP embedding and cell type annotations. The number of neighbours used to train the classifier was 20. The trained classifier was then applied to the query samples to get their predicted cell identities.

### Lineage emergence during blastocyst formation

To identify the emergence of Epi, Hyp and TE gene expression profiles during human blastocyst formation, we used the ESS to rank genes by how well their expression profiles overlap with the morula to Epi, Hyp or TE trajectories, and then progressively restricted the late blastocyst Epi, Hyp or TE populations. As an example, to identify Epi staging genes, we started by ranking genes based on their enrichment in the combined morula, ICM/TE branch, ICM and preImp-epi samples from our cell UMAP annotations (Fig 4A). This was followed by ranking against the combined ICM/TE branch, ICM and preImp-Epi population, and then by the combined ICM and preImp-Epi samples, and finally the preImp-Epi samples in isolation (Fig S7A). A similar process was repeated for the Hyp and TE trajectories. For each ranked gene list generated, we inspected the top 30 ranked genes and manually confirmed the gene expression profile using the UMAP embedding. Through this process we were able to identify sets of genes that delineate the emergence of Epi, Hyp and TE lineages from the morula progenitor population.

## Acknowledgements

We would like to thank Professor Marc Goodfellow, Professor Linus Schumacher and Dr. James Briscoe for their helpful discussions and insightful comments.

We thank Dr Dimitri Meistermann, Dr Cheng Zhao and Professor Fredrik Lanner for helping us obtain the spliced and unspliced scRNA-seq matrices used for RNA velocity analysis.

We thank Dr Manvendra Singh for providing cell label IDs for the Petropoulos et al. 2016 scRNA-seq data used in M. Singh et al. 2023.

Thank you to Dr Pengtao Liu and Dr Xi Chen for providing the human EPSCs scRNA-seq counts matrix used in Gao et al. 2019.

## Funding

This research was supported by the Medical Research Council Programme Grant (MR/W025310/1) and European Research Council Advanced Grant (Plastinet, contract 835312) funding to A.S.

A.S. is a Medical Research Council Professor (G1100526/2).

## Author contributions statement

Conceptualisation: A.R., A.S.; Data curation: A.R.; Formal Analysis: A.R.; Funding acquisition: A.S.; Investigation: A.R.; Methodology: A.R.; Project administration: A.S.; Resources: A.S.; Software: A.R.; Supervision: A.S.; Validation: A.R.; Visualisation: A.R.; Writing – original draft: A.R.; Writing – review & editing: A.R., A.S.

## Declaration of Interests

The authors declare no competing interests.

## 6 MATERIALS AND METHODS

### From discrete data Entropy Sorting to continuous data Entropy Sorting

A key distinction between the ESFW algorithm outlined in our previous work (A. Radley et al. 2023) and the cESFW algorithm presented in this work is that cESFW updates the theory behind ES so that it can be readily applied to continuous data. For a detailed explanation of how ES has been adapted for continuous data, see SI 1.

### A fifth entropy sorting error scenario

In addition to expanding the ES framework so that it may be applied to continuous data, we also introduce a fifth error scenario where ES ‘divergence can be observed through the ES parabola in a manner that indicates that samples may be displaying false negative (FN) expression levels. We present this new error scenario in Fig S8 (black box), alongside the 4 maximum entropy error scenarios introduced in our previous manuscript (A. Radley et al. 2023). The error scenarios are important because they provide implicit logic that determines how we calculate divergence and error potential for any pair of features.

As an example, in our newly proposed fifth error scenario, there are two features being compared against one another. The top feature (blue and red) is denoted the RF and the bottom feature (green and purple) is denoted the query feature (QF), because according to the maximum entropy principle, the QF should always be the feature with the larger cardinality for it’s observed minority state (A. Radley et al. 2023). For this pair of features, we see a strong but imperfect conditional relationship between the co-occurrence of ground truth (absent of X symbols) RF and QF minority states. However, the presence of RF majority states overlapping with QF minority states introduces uncertainty into the system, which manifests as a non-zero conditional entropy between the two features. Hence, we can hypothesise that the samples diverging from the ground truth (indicated by the blue X’s) are either real, or are present due to the introduction of erroneous data points. This allows us to set up a hypothesis test, which we introduce in A. Radley et al. 2023 as the ES hypothesis test. The ES hypothesis test allows us to quantify whether it is more likely or not than random chance that the overlapping RF majority and QF minority states are present due to sampling/measurement error.

All 5 of these error scenarios are encoded into our cESFW algorithm, which can be inspected to gain further insights into the algorithmic application of the error scenarios. However, prior to looking at our cESFW code, we advise reading the relevant error scenario sections in the supplemental of our A. Radley et al. 2023 publication for a detailed description.

### Continuous Entropy Sort Feature Weighting (cESFW) workflow

In Figure 1 we outlined the main steps in our proposed cESFW scRNA-seq data processing workflow. In this section we will provide further details for each step. Executable code to apply our cESFW workflow to the synthetic and human embryo scRNA-seq data presented in this paper can be found at https://github.com/aradley/cESFW_Embryo_Topology_Paper. The primary purpose of this section and our workflow is to give readers an insight into how cESFW can be successfully applied to scRNA-seq data. It is expected that users will find other ways to incorporate cESFW into their omics data analysis pipelines for the plethora of data that is now available to the community.

#### Temporary minority state normalisation

For features to be compared to one another with cESFW, they must each be converted into their minority state feature activity vectors (Fig S9A). Practically, converting features into their minority state vectors amounts to normalising each feature so that their values are within the same range, in a manner that meaningfully captures how features co-occur with one another. For gene expression data, this could be two genes where when one feature displays expression values close to its maximum expression in the data, the other gene also shows values close to its maximum expression. In this example, to convert these features to their minority state feature activity vectors, we could simply divide all the values of each feature by their respective feature maximums (Fig S9C). However, now consider that there is a third gene where when the first two genes have values close to 0, the third gene has values close to its maximum expression, and when the first two features have values close to their maximum, the third feature has values close to 0 (Fig S9A, B). This is equivalent to the high expression of the third gene being the majority state feature activity (Fig S9B), which is implicitly defined by the sum of activities in the third gene’s normalised vector being greater than half the number of samples. Hence, to convert the third gene to its minority state activity vector, we simply deduct each value of the vector from 1 (Fig S9A, B).

This logic is generally applicable to any dataset where we can justifiably convert each feature into active versus inactive states. How this abstraction is done and whether this is meaningful for different datasets is context dependent and ultimately left to the user. In this work, we found the relatively simple normalisation method presented in Fig. 7 is effective.

**Figure 7.**
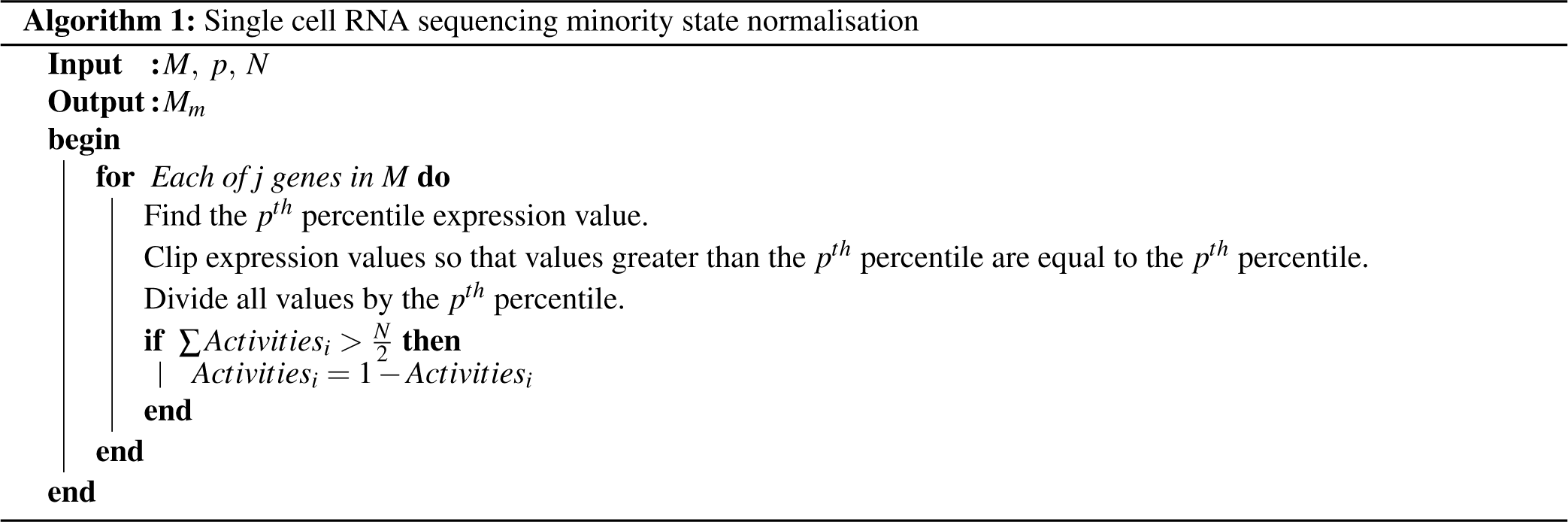
Where input *M* is the *i* by *j* raw scRNA-seq counts matrix (*i* = samples/cells, *j* = genes), *N* is the number of samples/cells comprising (*N* = *j*), and *p* is the percentile used to clip the maximum expression of each gene. In this work, *p* was set as the 97.5th percentile. The output, *M*_*m*_, is the normalised minority state activity *i* by *j* matrix.

Having obtained *M*_*m*_, we may now use it as the input for ES calculations to quantify the correlations between features.

#### Exclude genes significantly enriched within a dataset [Optional]

Sometimes when analysing scRNA-seq data we would like to combine datasets from different sequencing experiments. Combining datasets from different experiments can introduce batch effects as a significant confounder in differentiating biological signal from experimental noise. To mitigate the contribution of batch effects on downstream analysis of scRNA-seq data, several methods have been developed (Tran et al. 2020). Many of these methods focus on smoothing or regressing the counts matrices with the aim of making samples with similar cellular states that are derived from different datasets, look more similar to one another than samples of different cellular states that are from the same dataset. While there are scenarios where smoothing/imputation methods can be successfully employed to gain greater insights into scRNA-seq datasets, they always incur the risk of adding computational artefacts or unintentionally removing biological signal of interest.

In this cESFW workflow, we address the issue of batch effects in a different manner. Rather than change the values of the counts matrix, we hypothesised that genes that are significantly enriched in any individual dataset constitute genes that contribute to batch effects and hence should be removed from the data. In doing so, we aim to reduce batch effect signals to the point that biological patterns of interest that are common between datasets have a larger signal than dataset specific noise, which should in turn lead to more biologically relevant insights in downstream analysis.

To identify which genes are considered significantly enriched in a specific dataset in an unsupervised manner, we turn to the Error Potential (EP). The EP is a metric derived from Entropy Sorting and described in detail by A. Radley et al. 2023. Given two minority state activity feature vectors, the EP between the two features can be calculated, and if the EP > 0, the features can be considered to be co-occurring with each other in a manner that is greater than random chance. In Fig 8 we provide pseudocode for how we use the EP for excluding features that are significantly enriched in any of the datasets comprising a scRNA-seq counts matrix of interest.

**Figure 8.**
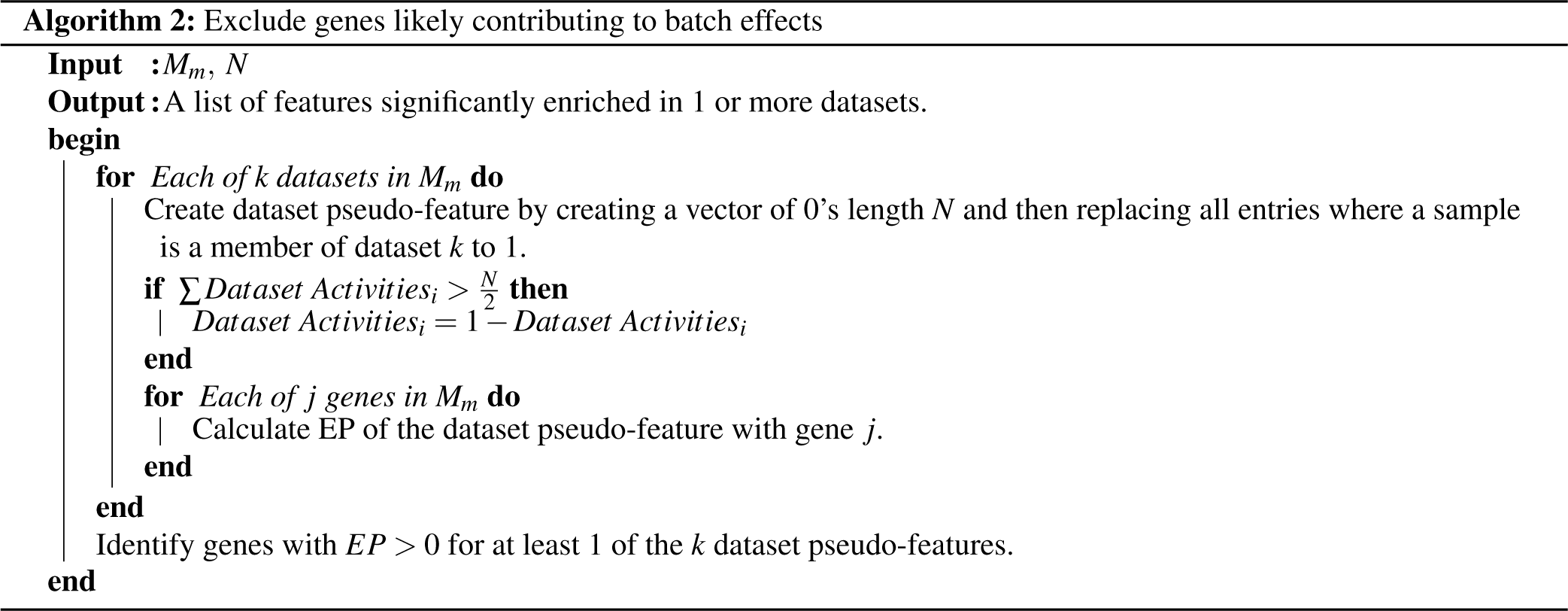
Where input *M*_*m*_ is the *i* by *j* normalised minority state activity (*i* = samples/cells, *j* = genes), *N* is the number of samples/cells comprising (*N* = *j*) and *k* is the number of datasets the were combined to create *M*_*m*_.

A limitation with this batch specific gene exclusion methodology is that if there are datasets within the combined counts matrix (*M*_*m*_) that are comprised of almost entirely one cell type, and that cell type has a distinct gene signature that is not present in any other dataset, these genes may be unwantingly identified as batch effect genes and removed from the data. In such a scenario, it would be up to the researcher to modify the workflow accordingly or seek alternative methodologies.

#### Calculate ESS and EP feature matrices

Having created the *M*_*m*_, we use it to calculate the Entropy Sort Score (ESS) and Error Potential (EP) pairwise for each gene in *M*_*m*_. In doing so we generate the Entropy Sort Scores (*ESSs*) and Error Potentials (*EPs*) matrices, which are *j* by *j* matricies where *j* is the number of genes present in *M*_*m*_. The ESS uses ES theory to quantify the correlation between two features with a value between -1 and 1. The EP uses a hypothesis test between the prospect of two features being completely independent versus the two features being dependent, to quantify the significance of the correlation between the two features. When EP < 0, it is more likely that the observed correlation between two features occurred through a random relationship between two independent features than the features having a dependent relationship. Conversely, when EP > 0, the correlation between the two features is more likely to have been observed due to some dependent relationship.

For further details regarding the theory and derivation behind the ESS and EP, see A. Radley et al. 2023. In the next steps of our cESFW workflow, we use the *ESSs* and *EPs* matrices to get a feature importance weight for each of the *j* genes, and use these weights to decide which genes should be retained in the scRNA-seq counts matrix, and which should be excluded.

#### Calculate cESFW feature weights

Having generated the *ESSs* and *EPs* matrices, we now calculate the cESFW feature weights for each of the *j* genes. To calculate gene weights, we combine the *ESSs* and *EPs* matrices derived from ES theory, with the concept of node centrality from graph theory. In the context of scRNA-seq data, node centrality can be used to try to distinguish between genes (nodes) that form networks with relatively high correlations with other genes in the dataset, from genes that form weak correlative networks. The assumption is that genes that are members of highly correlating networks are more likely to be involved in cellular function/identity.

Because the ESS is a symmetric correlation metric, the *ESSs* matrix can be thought of as a weighted undirected graph in the same manner as other correlation metrics (Pearson’s correlation, mutual information, etc.), where *ESSs*[2, 4] = *ESSs*[4, 2] = 0.7 would quantify the weight of the edge between genes 2 and 4 as 0.7. However, practical limitations of node centrality often force users to apply supervision to the analysis of complex datasets such as scRNA-seq data. For example, A. Singh, R. R. Singh, and Iyengar 2020 discuss how node centrality measures that use weighted edges can fail to account for the number of edges that each gene has to other genes. In the following we show that with the *ESSs*_*jx j*_ and *EPs*_*jx j*_ matrices we can calculate node centralities of weighted networks in a manner that is unsupervised and accounts for the number of edges that each gene has to other genes.

As stated, *ESSs* can be used as the weighted graph. Now note that each entry of *EPs* quantifies to what degree the corresponding correlation value in *ESSs*_*jx j*_ is likely to have occurred by random chance or a dependent relationship between a pair of features. According to ES hypothesis testing (A. Radley et al. 2023), whenever EP < 0, the feature pair correlation can be assumed to have occurred by random chance. We utilise this property to create a dependent *EPs* matrix, *dEPs*, where *dEPs* is created by setting all values in *EPs* < 0 to 0. *dEPs* now quantifies all the pairwise gene relationships whose co-expression occurs to a degree greater than random chance. Now we can calculate the weighted node centrality of each gene as the weighted column averages of *ESSs* with *dEPs* as weights such that;

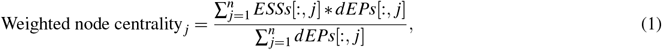

where *j* denotes the column/gene index in *ESSs* and *dEPs*. By weighting each value in *ESSs* by the corresponding values in *dEPs*, we mitigate the contribution of uninformative/random correlative patterns to the calculation of gene centrality.

Finally, we encode a biologically inspired assumption into the Weighted node centrality *j* to get our final cESFW gene weights. Because every non-zero value in *dEPs* indicates that a pair of genes have a quantifiably significant relationship, we can count the number of quantifiably meaningful edges a gene has with other genes in the data. We then seek to penalise genes that have significantly more edges than others through the following;

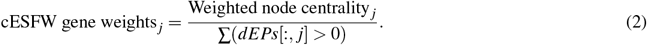

The biological assumption behind Eqn 2 is that cell identity and differentiation is controlled by relatively small networks of tightly controlled genes. Hence, genes with relatively high weighted node centralities (Eqn. 1) and relatively low numbers of significant gene co-regulation edges, should be more informative of cellular identity. Conversely, genes that have relatively low weighted node centralities and/or relatively high numbers of significant gene co-regulation edges should constitute genes that are weakly or non-informative of cell identity.

In summary, we have calculated a vector of cESFW gene weights, for each gene in *ESSs*_*jx j*_, such that a larger cESFW gene weight indicates that a gene’s expression in the original scRNA-seq counts matrix are more likely to be part of a small network of highly correlated genes than other genes with lower cESFW gene weights. As such, we can use the cESFW gene weights to form a ranked list of genes for downstream analysis.

#### Select gene cluster with branching network

The final step of our proposed cESFW workflow is to take the cESFW gene weights and use them to select the top most informative genes for downstream analysis. In a similar manner to HVG analysis, we can simply take a subset of the top highest ranking genes according to the cESFW gene weights. Up until this point, the cESFW workflow has been entirely unsupervised, i.e., no subjective user decisions were applied, and no prior knowledge regarding cell type or gene importance was used when deciding which genes to exclude from downstream analysis. As with HVG analysis, the final step of the cESFW workflow where we choose how many genes to select based on the cESFW gene weights is an iterative process where researchers use domain knowledge to settle on the final gene set, based on the assumption that the cESFW ranked gene list provides a meaningful proxy for thresholding. Below we provide a description of our rational for using the cESFW gene weights to identify a set of biologically informative genes for the human embryo scRNA-seq data within this manuscript. We present this description as a guide to aid analysis on other datasets.

For the human embryo scRNA-seq data presented in this manuscript, we selected the top 4000 genes by taking the 4000 genes with the highest cESFW gene weights. The selection threshold of 4000 was chosen by looking at UMAPs of the genes in the ESSs matrix at varying cESFW gene weight cutoffs (Fig S10A). In these UMAPs we qualitatively look for two main properties. Firstly we are looking for a clusters of genes that have significant branching patterns, rather than smooth “blob” like clusters. In Fig S10A we see that as we decease the number of top ranked genes that we include, the darker blue coloured cluster forms increasingly distinct branches. These branches indicate that there are genes that form fully connected networks, with some genes being very specifically connected to each other without being directly connected to many other genes in the network. Biologically we infer that these represent genes that are specifically expressed in a subset of cells present in the entire dataset, and are hence genes of interest when dissecting cellular identity. The top 4000 genes was chosen relatively arbitrarily as the point at which a clear branching pattern appears in the dark blue cluster. Readers will notice that choosing values smaller than 4000 can result in even more distinct branches in the blue cluster. We suggest users of our cESFW workflow aim to find a balance between keeping as many genes as possible while increasing the qualitative property of branching structure.

The second feature of these UMAPs that we are interested in is that a separate cluster of genes with significantly higher weighted node centrality scores can appear (yellow). Empirically we find that failure to remove these clusters of genes prevents the emergence of high resolution scRNA-seq embeddings (see the online GitHub workflow for examples). Gene set enrichment analysis of the final 3012 gene set used throughout this manuscript, comprising the blue cluster of the top 4000 genes in (Fig S10A, red square), showed that these genes were enriched for gene ontology (GO) terms relating to regulation and development (Fig S10B). Conversely, the GO terms of the 988 genes comprising the yellow cluster were dominated by terms relating to transcription, translation and metabolism (Fig S10C). Hence we hypothesise that the genes present in the relatively highly correlated yellow clusters represent expression signals relating to generic cell transcription, translation and metabolism processes which are less discriminating of cellular identity/function than genes that are part of the blue branching cluster. By excluding the genes from the yellow cluster in downstream analysis, we appear to further increase the signal to noise ratio of cell type specific expression, which ultimately improves our ability to discriminate different cell identities in the data.

We emphasise that this final step of the cESFW workflow is supervised and subject user interpretation/dataset variability. We hope that our transparency regarding this step helps guide potential users of our cESFW workflow to improved resolution in their datasets, or to develop new cESFW feature weight based selection criteria themselves.

## MAIN FIGURE LEGENDS

To be filled in later for submission.

## SUPPLEMENTARY MATERIAL

### Supplemental Figures

**Table S1.** A placeholder table for annotated cell type ranked gene list table which is ready to be uploaded for submission.

| Col1 | Col2 | Col2 | Col3 |
| --- | --- | --- | --- |
| 1 | 6 | 87837 | 787 |
| 2 | 7 | 78 | 5415 |
| 3 | 545 | 778 | 7507 |
| 4 | 545 | 18744 | 7560 |
| 5 | 88 | 788 | 6344 |

**Figure S1.**
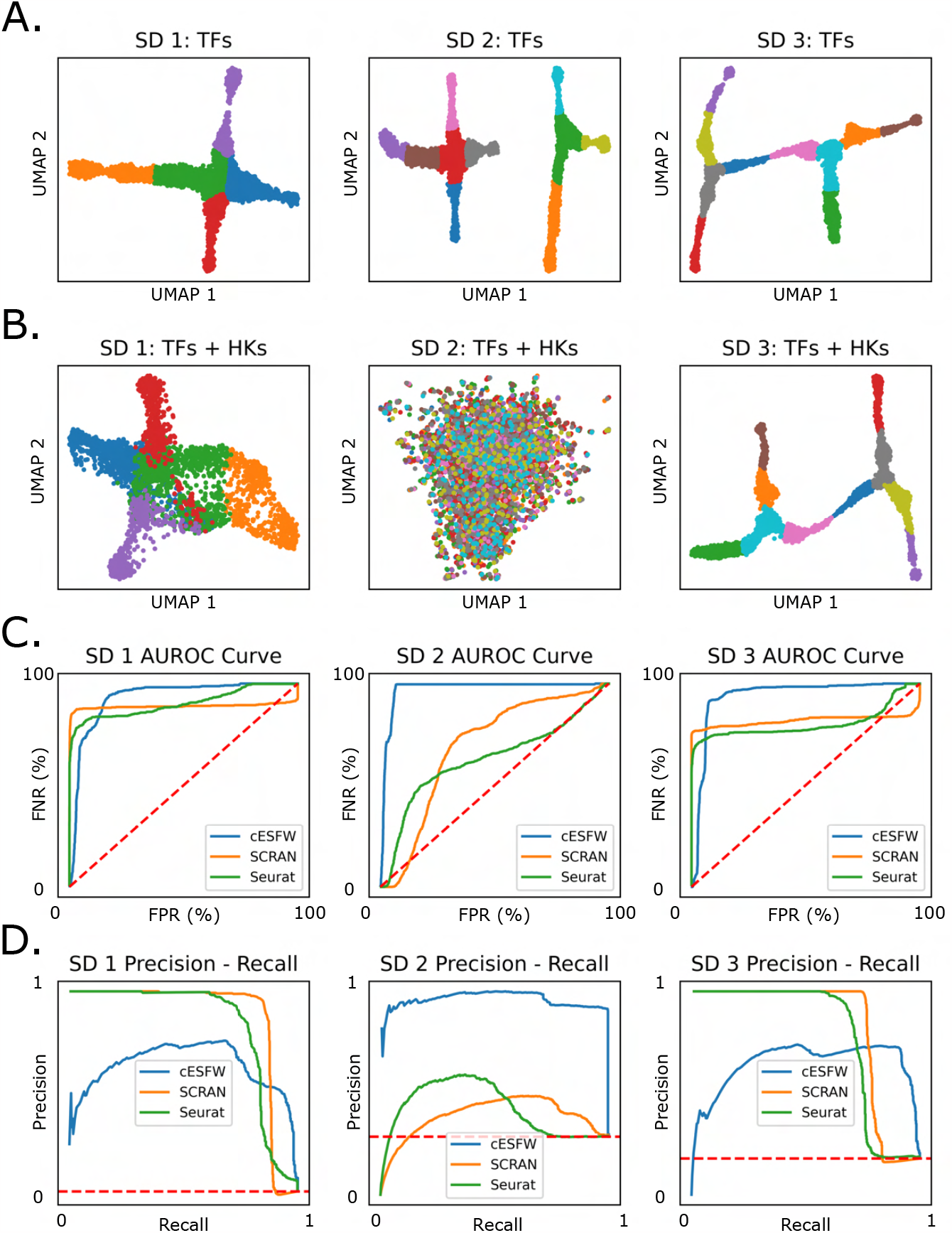
Dyngen generated synthetic datasets (SDs) 1-3. **A.** UMAPs of SDs generated using just the TFs of each dataset. **B**. UMAPs of SDs generated using all of the TF and HK genes of each dataset. **C**. AUROC curves generated by different feature selection methods on each of SDs 1-3. **D**. PR-AUCs generated by different feature selection methods on each of SDs 1-3.

**Figure S2.**
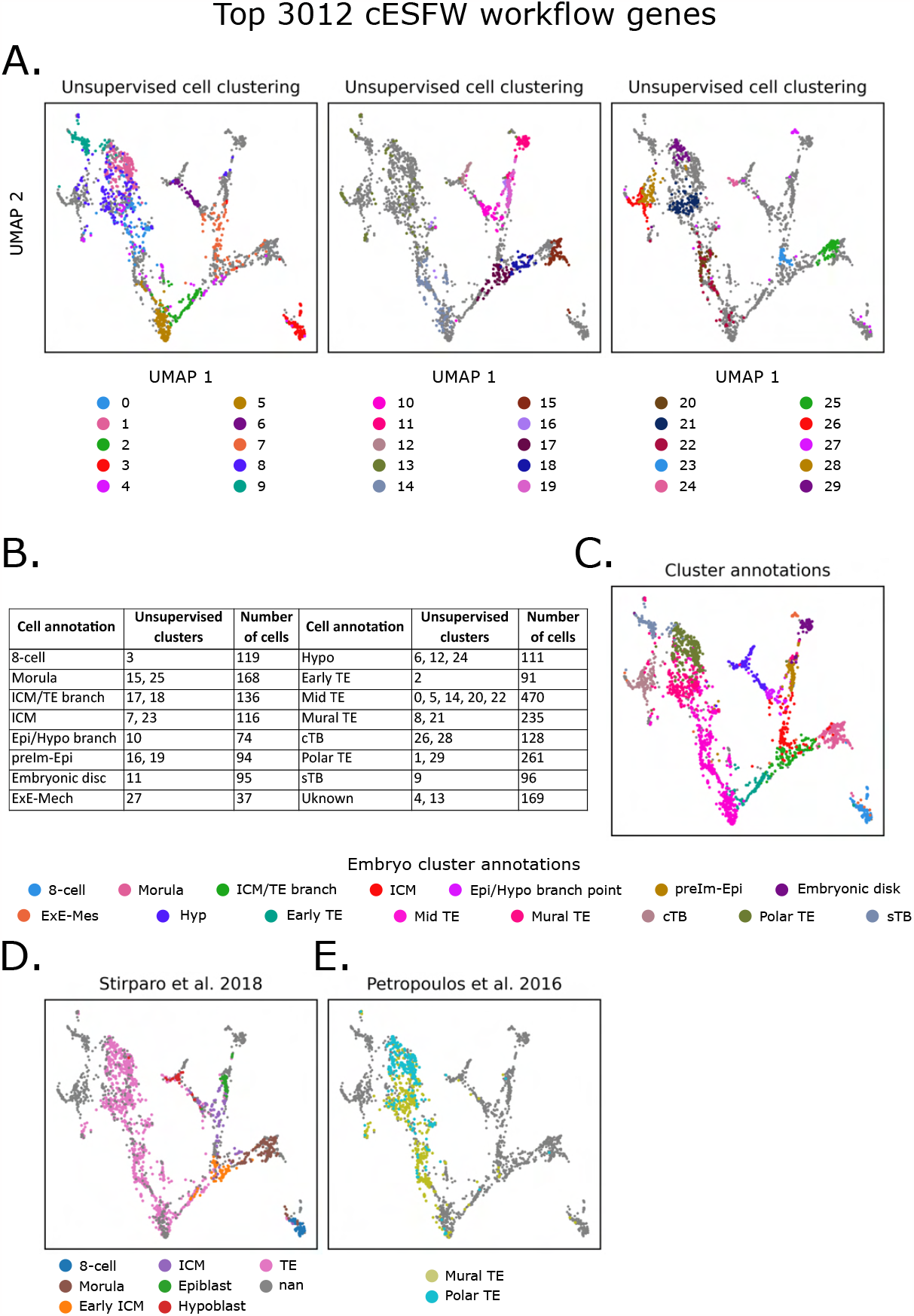
Annotation of our UMAP embedding. Unsupervised agglomerative clustering with number of clusters set to 30. Clustering was performed on the raw counts matrix subsetted down to our 3012 cESFW selected genes. **A**. The resulting sample clusters visualised on our UMAP embedding. By inspecting known markers and previous analyses of these scRNA-seq data, we manually groups our unsupervised clusters into annotated cell types. **B**. Summary of the annotated cell types and which unsupervised clusters were used to form them. **C**. Visualisation our our annotated cell types on our UMAP embedding. **D**. Stirparo et al. 2018‘s supervised analysis of the Petropoulos et al. 2016 scRNA-seq data support our cell type annotations. **E**. Petropoulos et al. 2016‘s supervised analysis for differentiating between mural and polar TE support our cell type annotations.

**Figure S3.**
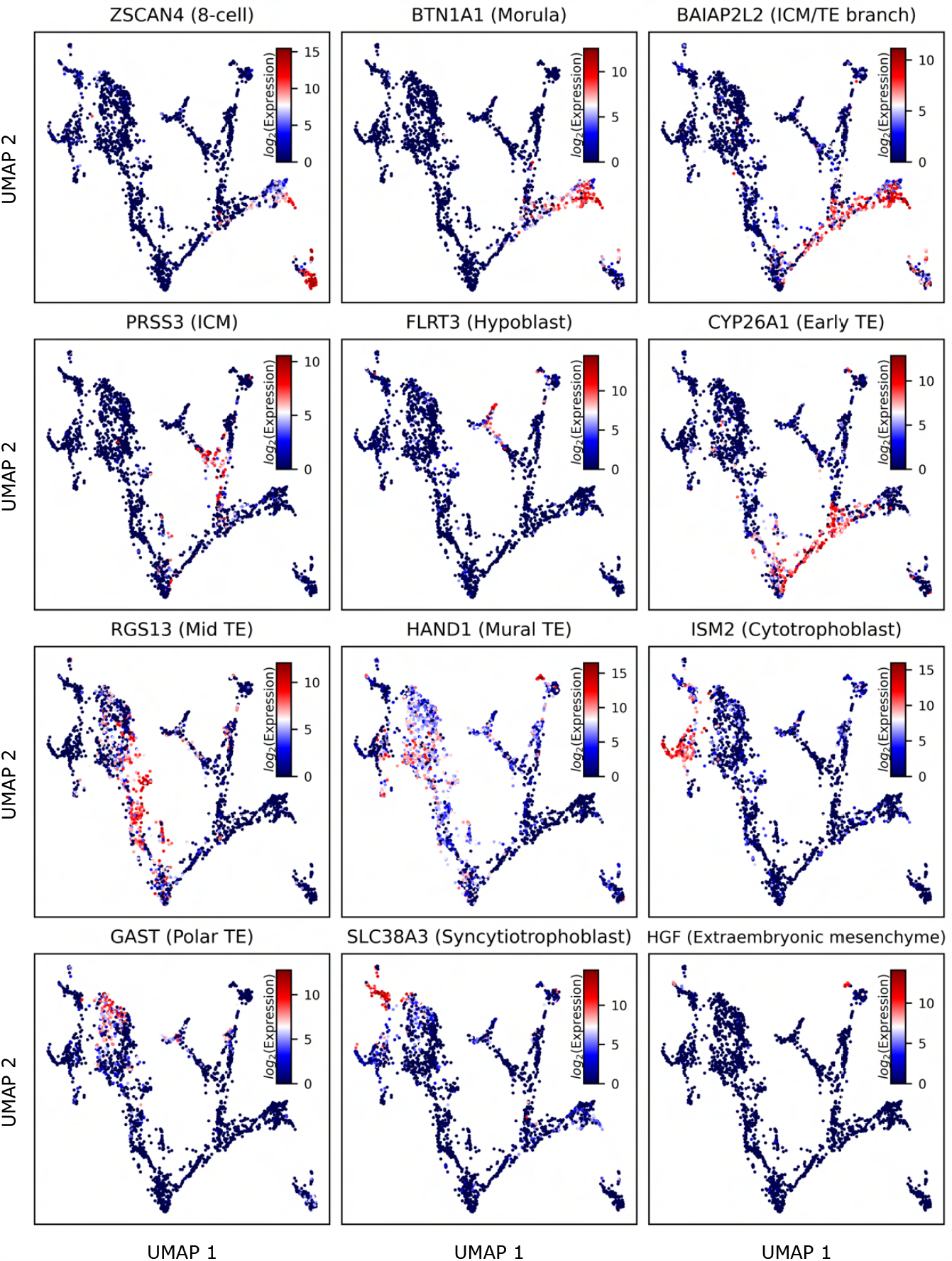
Visualisation of cell type markers supported by the literature. Marker gene references are as follow: **8-cell** - Taubenschmid-Stowers et al. 2022; **Morula** - Stirparo et al. 2018; **ICM/TE branch** - Singh et al. 2023; **ICM** - A. Radley et al. 2023; **Hypoblast** - Corujo-Simon, A. H. Radley, and Nichols 2023; **Early TE** - Liu et al. 2022; **Mid TE** - Zadora et al. 2017; **Mural TE** - Liu et al. 2022; **Cytotrophoblast** - Li, Kurosawa, and Iwata 2019; **Polar TE** - Yue et al. 2020; **Syncytiotrophoblast** - Yabe et al. 2016; **Extraembryonic mesenchyme** - Yang et al. 2021.

**Figure S4.**
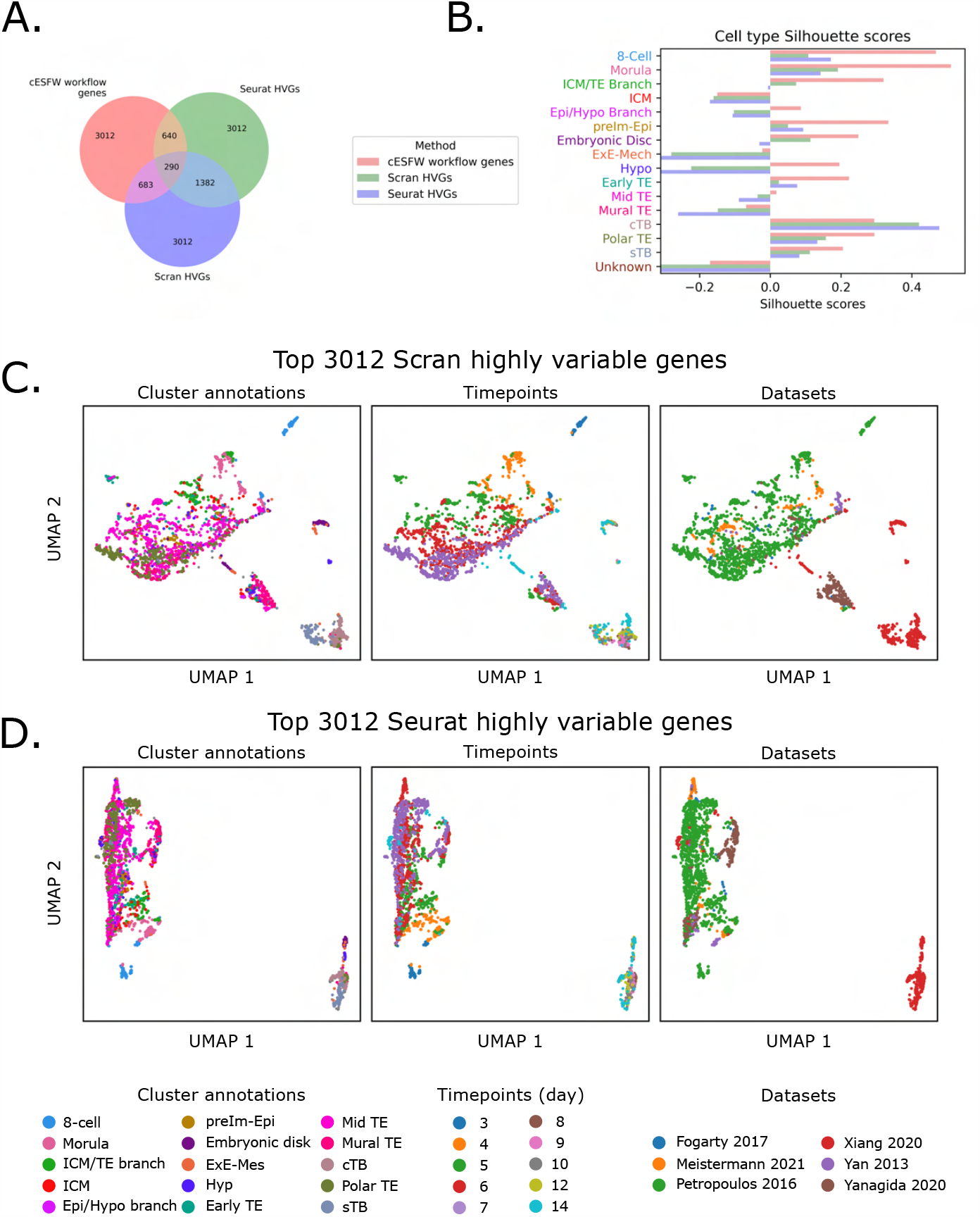
Comparison 3012 cESFW workflow genes against top 3012 Scran and Seurat HVGs. **A.** Venn diagram of overlap between the genes within each of the 3012 gene sets. **B**. Silhouette scores of each of our annotated clusters when using each of the 3012 gene sets. **B, C**. UMAP generated when using the same scRNA-seq counts matrix and UMAP parameters used to generate our cESFW UMAP embedding, but instead using the top 3012 Scran HVGs (B.) or top 3012 Seurat HVGs (C.).

**Figure S5.**
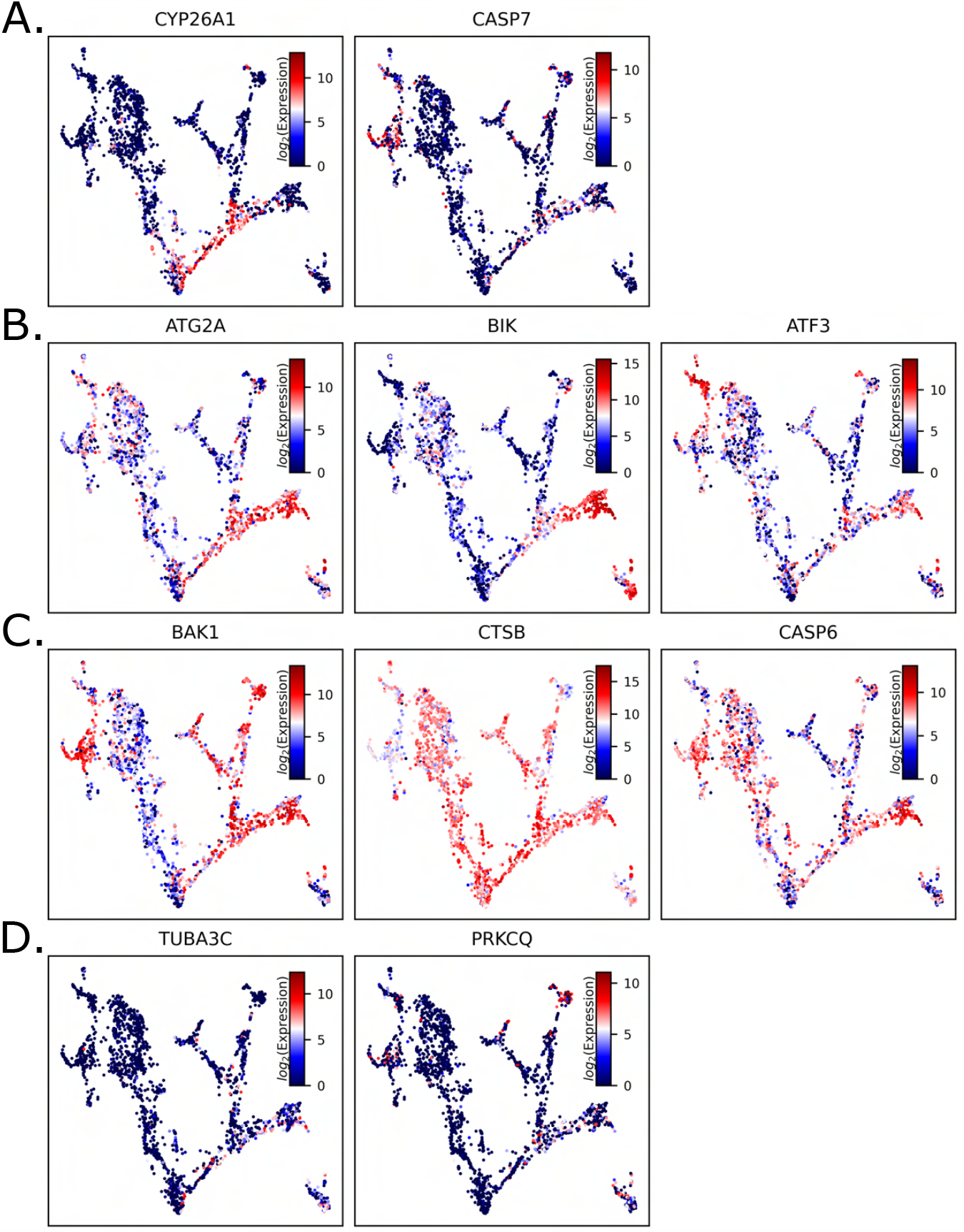
Proposed apoptotic markers of NCC cells display NCC non-specific expression profiles. Visualisation of NCC specific apoptosis markers proposed by Singh et al. 2023 show a variety of NCC specific and non-specific expression patters. **A**. Proposed apoptosis markers that have relatively specific NCC or ICM/TE branch expression. **C**. Proposed apoptosis markers that show higher expression in morula and/or 8-cell populations than NCC or ICM/TE cells. **C**. Proposed apoptosis markers that are upregulated across several cell types of the day 3-14 human embryo. **D**. Proposed apoptosis markers that show sparse/low expression throughout the day 3-14 human embryo.

**Figure S6.**
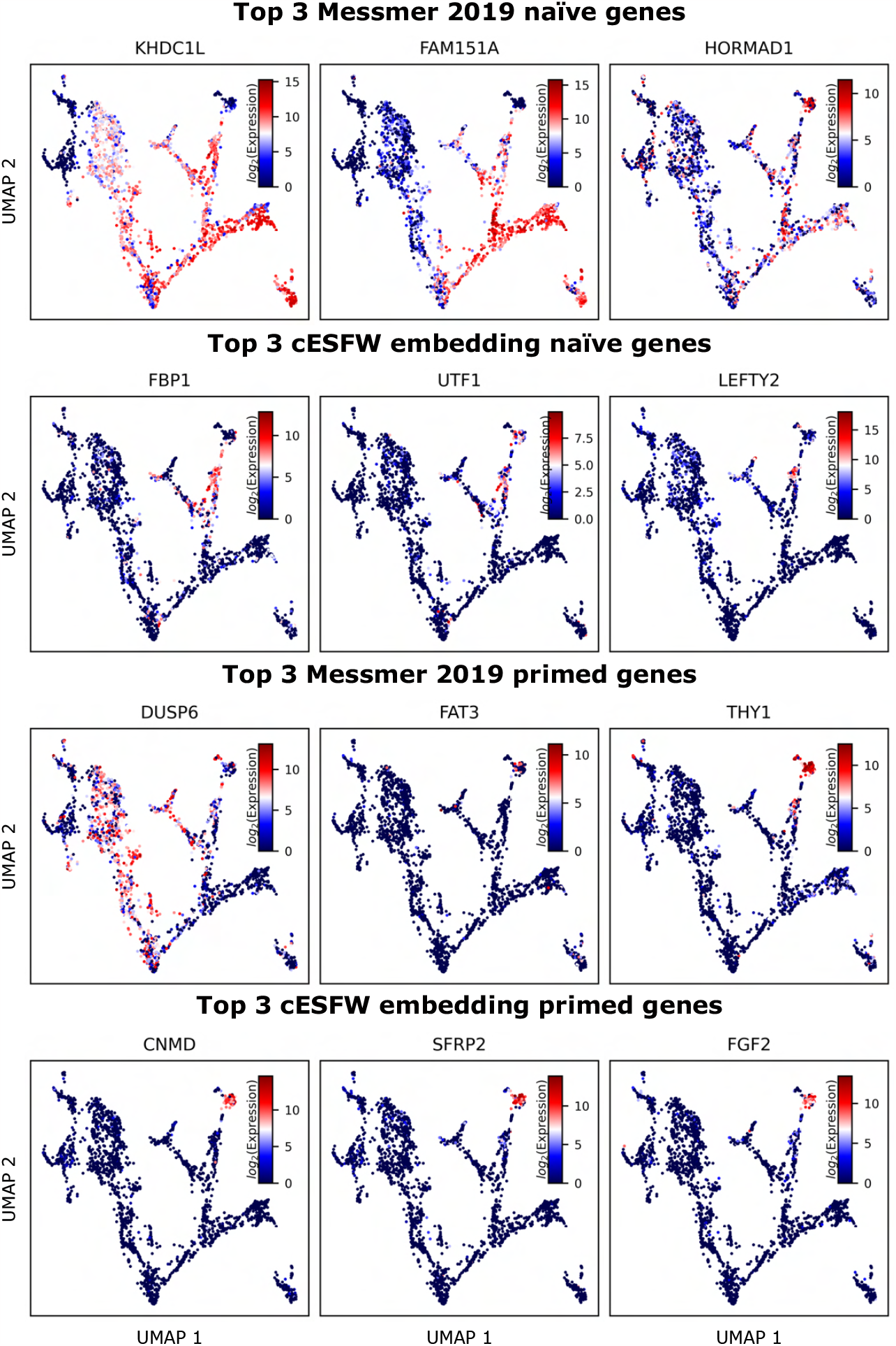
Comparison of top 3 Messmer et al. 2019 naïve and primed genes against the top 3 naïve and primed genes according to our cell type annotation ranked gene lists (Table S1).

**Figure S7.**
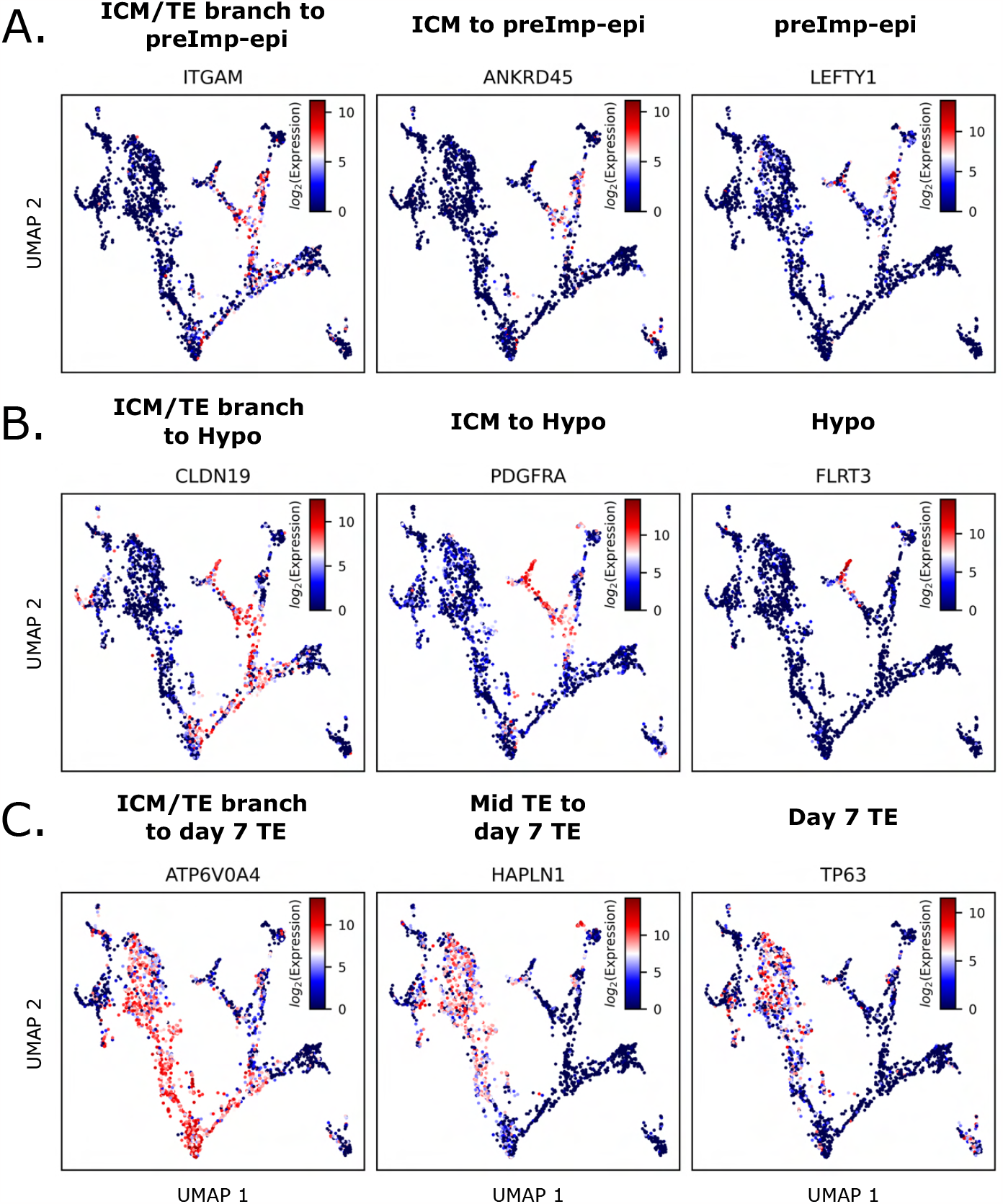
Emerging epiblast, hypoblast and trophectoderm signatures during blastocyst development. UMAP gene expression profiles for example genes of the Epi (A.), Hypo (B.) or TE (C.) lineages. Presented genes are from those that are highlighted in green in the heatmaps shown in Fig 6.

**Figure S8.**
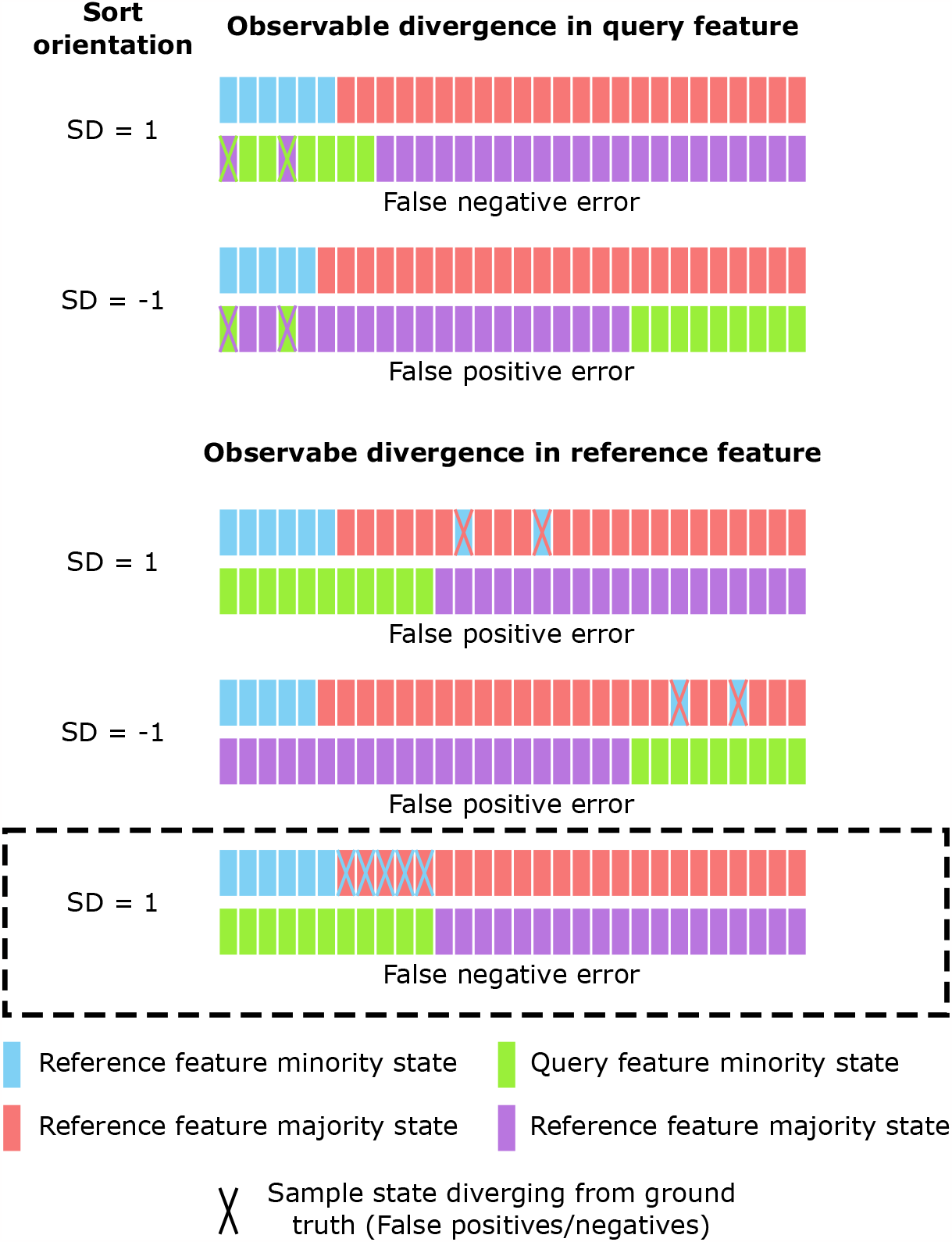
Entropy sorting error scenarios. A set of simple examples that demonstrate the 5 known error scenarios where expression states between two features can be quantified as potential false positive (FP) or false negative (FN) data points through the ES parabola. SD indicates the split direction for the observed reference feature (RF) and query feature (QF) pair. These examples are set up with discrete states (minority Vs majority) for simplicity, but can be expanded to continuous data, as per SI 1. The top 4 scenarios were outlined in detail in our previous work (A. Radley et al. 2023). The bottom error scenario (black box) outlines a previously undescribed error scenario where ES divergence indicative of FN expression values can be observed.

**Figure S9.**
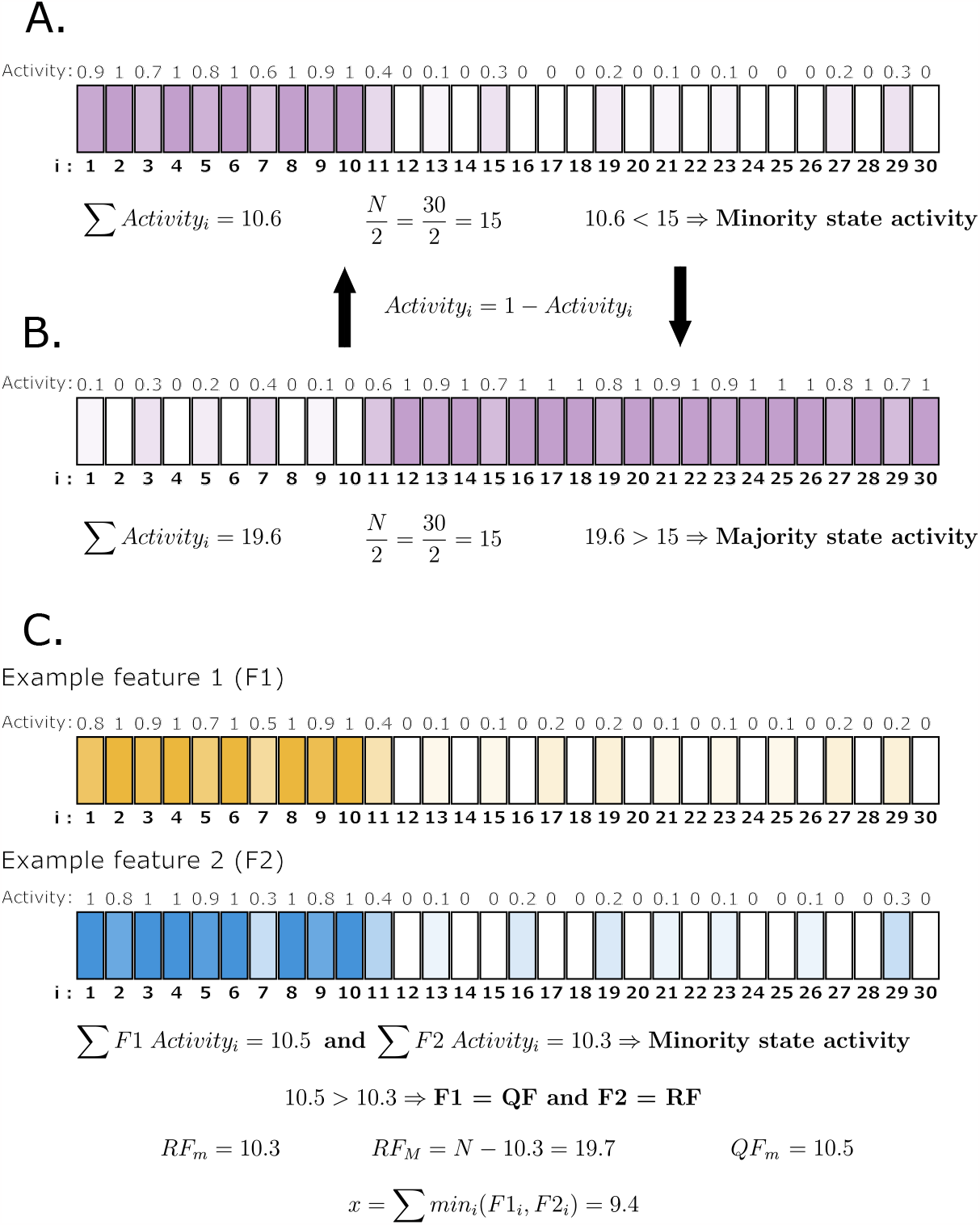
Calculating the inputs of the ESE for continuous data. **A, B.** An example feature where the sample activities are represented in gradients of purple. In A. the feature is being observed in a form that represents its minority state activities, since the sum of the activities for all 30 samples is less than ^*N*^ = 15. We can take the same feature and view it in its majority state activities by deducting the minority state activities of each sample from 1, as shown in B. The majority state form in B. can then be returned to the minority state form in A. by the same process. **C**. A pair of features in the minority state activity forms. Since ∑ *F*1*Activity*_*i*_ > ? *F*2*Activity*_*i*_, F1 is the QF. Following this, we can find *RFm, RF*_*M*_, *QF*_*m*_ and *x*, so that the CE between F1 and F2 can be calculated via the ESE (Eqn 2).

**Figure S10.**
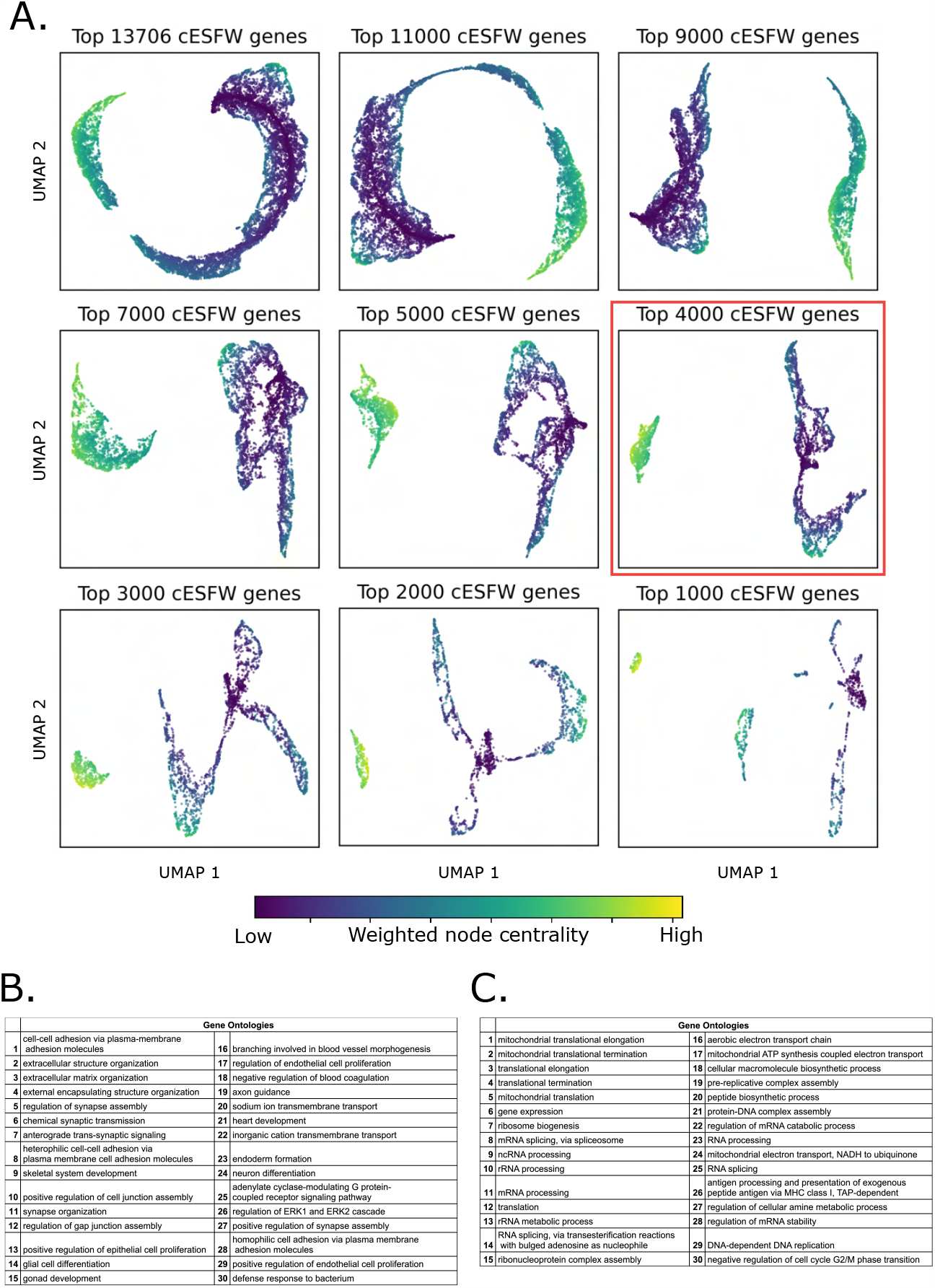
Identifying a cluster of genes with branching structure. **A.** UMAPs of top ranked genes with increasingly stringent thresholds shows the emergence of a cluster of genes that form a graph with branches. We hypothesise that these branches indicate sets of co-regulating genes. To generate the high resolution human embryo embedding show in the main text of this paper, we used a cESFW selection threshold that took the top 4000 genes according to cESFW weighting (red box in this figure). **B**. Gene ontology analysis of the genes in blue branching cluster of the top 4000 genes (red box) shows that these genes are enriched for developmental and differentiation terms. **C**. Gene ontology analysis of the genes in yellow cluster of the top 4000 genes (red box) shows that these genes are enriched for cell transcription, translation and metabolic processes.

## Supplementary Information

### SI 1 From discrete data Entropy Sorting to continuous data Entropy Sorting

Entropy Sorting (ES) is a mathematical framework that quantifies the correlations between features by reformulating conditional entropy as a sorting problem, rather than a probabilistic one. By imagining the relationships between features as sorting problems, ES gives us access to mathematical properties that are particularly useful when analysing high dimensional data. In A. Radley et al. 2023, the authors comprehensively outline the derivation of ES and demonstrate how properties of ES can be helpful when interrogating high dimensional data. The foundation of ES is the Entropy Sort Equation (ESE), which given two features, calculates their conditional entropy (CE). (A. Radley et al. 2023) defined the ESE as;

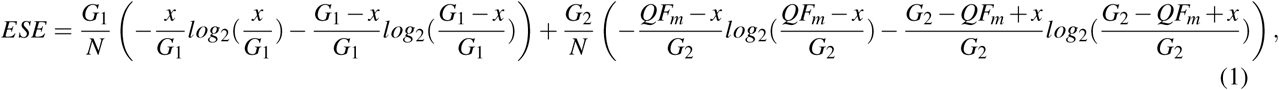

where *RF* and *QF* stand for the Reference Feature and Query Feature respectively, and *N* equals the number of samples that comprise the RF/QF. According to the maximum entropy principle, given two features the *QF* is designated as the feature with the higher independent entropy, and as such the *RF* is the feature with the lower independent entropy. A. Radley et al. 2023 derive the ESE from a discrete state scenario where a feature can only be observed in one of two states, i.e. 0 or 1, where 1 is always designated as the minority (m) state. For example if *N* = 30 and we observe 20 samples of a feature to display 1’s and 10 to display 0’s, we must switch the 0 and 1 values for each sample such that there are instead 10 1’s and 20 0’s. Following this, *G*_1_ is the number of samples in the *RF* that are equal to 1 and *G*_2_ equals the number of samples in the *RF* equal to 0. Hence, *G*_1_ + *G*_2_ = *N*. Likewise, *QFm* is equal to the number of samples in the *QF* that display the minority states (i.e. are equal to 1). Finally, *x* is the only variable in the ESE, and denotes the number of samples where samples both the *RF* and *QF* are equal to 1. We call *x* the overlap between the *RF* and *QF* minority states.

An intuitive way of interpreting the ESE is that it quantifies the degree to which the less common observation of two features overlap with one another. In the context of gene expression data, samples displaying 0 could be when a gene is inactive in a cell and those displaying 1 have the gene as active. The ESE then quantifies how well the active states of both genes co-occur with one another. A clear limitation of the above ESE formulation is that many datasets, including gene expression data, consist more than two states/values. Here we demonstrate how a simple alteration to the ES framework allows us to expand the usage of ES to datasets with continuous values.

We start by noting that rather than calling the 0 and 1 values “states”, we can refer to them as activities, where 0 indicates the feature is completely inactive and 1 designates the the feature is fully active. The notion of activity facilitates an interpretable meaning of a sample displaying a value of 0.5 for a feature, which would mean that the feature is 50% active in the given sample. In the MATERIALS AND METHODS we discuss how we ensure all values in a scRNA-seq dataset are placed with a range of 0-1. For this section, we will use the examples in Figure S9 to redefine each of the terms in the ESE to be conducive with continuous data in the following manner:

- *N*: Remains as the number of samples comprising the *RF*/*QF*.
- _*m*_: Minority state activity. In the discrete case, this was the number of samples displaying the minority state. In the continuous case, _*m*_ is equal to the sum of all minority state activities. In Figure S9A, _*m*_ = 10.6.
- _*M*_: Majority state activity. In the discrete case, this was the number of samples displaying the majority state. In the continuous case, _*M*_ is equal to the sum of all majority state activities. In Figure S9B, _*M*_ = 19.4.
- *QF*: Of the two features being compared against one another, the QF is designated as the feature with the larger minority state activity (_*m*_). Following this definition, in Figure S9C the QF is F1.
- *RF*: For a pair of features being compared against one another, the RF is designated as the feature with the smaller minority state activity (_*m*_). Following this definition, in Figure S9C the RF is F2. In cases where F1 and F2 have equal cardinalities, one should arbitrarily assign a feature as the RF/QF.
- *G*_1_: In the ESE derived by (A. Radley et al. 2023) (Eqn (1)), *G*_1_ is equivalent to *RF*_*m*_, hence to make nomenclature more consistent, from now on we substitute *RF*_*m*_ in place of *G*_1_. In Figure S9C, *RF*_*m*_ = 10.3.
- *G*_2_: In the ESE derived by (A. Radley et al. 2023) (Eqn (1)), *G*_2_ is equivalent to *RF*_*M*_, hence to make nomenclature more consistent, from now on we substitute *RF*_*M*_ in place of *G*_2_. In Figure S9C, *RF*_*M*_ = 19.7.
- *QF*_*m*_: The QF minority state activity. In Figure S9C, *QF*_*m*_ = 10.5.
- *x*: The minority state activity overlap. In the continuous case, the overlap may be thought of as the sum of the minority state activities that co-occur in individual samples for both features. Mathematically, this is the minimum value observed in each sample for both features. In Figure S9C, *x* = 9.4.

Substituting these new continuous form definitions into the ESE leaves us with the following equation:

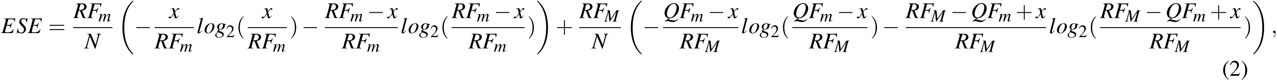

which only varies from Eqn 1 due to the nomenclature substitution of *RF*_*m*_ and *RF*_*M*_ for *G*_1_ and *G*_2_ respectively. Hence we have demonstrated that the above definitions leading to Eqn 2 allow us to apply the principles of ES to continuous data.

## Notes

### Competing Interest Statement

The authors have declared no competing interest.

### Summary of Updates

1) Amended a typo in the abstract. 2) Added the supplemental table to the submission. The table was referenced in the original submission, but was not uploaded to the public domain.

